# Genomic splice-donor disruption rescues systemic disease in progeria

**DOI:** 10.64898/2026.09.17.752274

**Authors:** Wei Jin, Jingchun Ma, Hao Yuan, Yu Zou, Zixuan Zhang, Jinhang Zuo, Sheung Kin Ken Wong, Tuo Li, Xiaomei Lu, Zhongjun Zhou

**Affiliations:** School of Biomedical Sciences, LKS Faculty of Medicine, The University of Hong Kong, Hong Kong; Dongguan Institute of Pediatrics, Dongguan Children’s Hospital, Dongguan, China; Department of Computer Science, City University of Hong Kong, Hong Kong; Department of Endocrinology, Changzheng Hospital, Shanghai, China; Guangdong Provincial Key Laboratory of Medical Immunology and Molecular Diagnostics; Institute of Aging Research, Guangdong Medical University, China

**Author notes:** These authors contributed equally (Wei Jin, Jingchun Ma). Correspondence should be addressed to Wei Jin or Zhongjun Zhou.

## Abstract

RNA splicing determines which protein isoforms a gene produces, and aberrant splice-site activation can cause severe disease^1,2^. In Hutchinson–Gilford progeria syndrome, a synonymous *LMNA* mutation strengthens a cryptic 5′ splice donor site to produce progerin, a toxic lamin A isoform driving premature cardiovascular death^3,4^. Within this cryptic donor, the activating mutation lies near the canonical GT dinucleotide required for splice-site recognition. A single guide can therefore preferentially pair with the mutant sequence while positioning Cas9 cleavage at this dinucleotide. Here we show that CRISPR/Cas9-mediated disruption of this splice site suppresses progerin without requiring precise correction of the activating mutation and prolongs lifespan in progeroid mice. In patient-derived fibroblasts and induced pluripotent stem cell-derived mesenchymal stem cells, this intervention reduced progerin by 92.7–96.8% while retaining lamin A and lamin C expression. A single neonatal injection of AAV9 carrying the corresponding mouse guide increased median survival from 115 to 279 days in Cas9-expressing *Lmna*^G609G/G609G^ mice. Treatment attenuated aortic, skeletal and multiorgan pathology. Treated homozygous pairs produced 52 offspring across seven litters. These findings establish pathogenic splice signals as genomic targets for sustained suppression of toxic isoforms while retaining canonical gene output.

## Main Text

Pre-mRNA splicing is a central determinant of transcript identity, and altered splice-site selection is a recurrent mechanism of inherited and acquired diseases^1^. Cryptic donor or acceptor sequences that are normally suppressed by the surrounding sequence context can be unmasked by single-nucleotide changes, including synonymous substitutions, redirecting a fraction of transcripts towards aberrant isoforms^2,5^. Such variants contribute to β-thalassaemia^6^, cystic fibrosis^7^, spinal muscular atrophy^8^, and familial hypercholesterolemia^9^ and define cancer subgroups, including *MET* exon 14-skipping non-small-cell lung cancer^10,11^.

Hutchinson–Gilford progeria syndrome (HGPS) is caused by a *de novo* synonymous substitution in *LMNA* (c.1824C>T; p. Gly608Gly) that strengthens a cryptic 5′ splice donor in exon 11, diverting a proportion of transcripts to an aberrant junction that removes 150 nucleotides and produces progerin^3,12^. Progerin retains a farnesylated C-terminal tail, accumulates at the nuclear envelope and disrupts lamina integrity, heterochromatin organisation and DNA repair, leading to progressive multi-organ deterioration and premature cardiovascular death^4,13^.

Existing HGPS therapies intervene at different levels of the progerin pathogenic cascade. Farnesyltransferase inhibition reduces progerin toxicity at the protein level^13^. Antisense oligonucleotides (ASOs) inhibit pathogenic splicing^14^ or redirect *LMNA* processing towards lamin C^15^ at the RNA level, requiring sustained dosing. CRISPR/Cas9-mediated indels in the lamin A/progerin coding region ablate progerin together with lamin A^16,17^. Adenine base editing reverts the causative nucleotide itself^18^. These strategies demonstrate that progerin is therapeutically tractable at the levels of post-translational modification, transcript output, and mutation correction. Whether the pathogenic splice signal itself can be disabled durably at the DNA level to prevent toxic-isoform production while retaining canonical isoform expression remains unexplored.

Here we developed a CRISPR/Cas9-editing based progerin splice-site silencing (PSS) strategy, in which Cas9 cleavage is directed to the +1/+2 GT dinucleotide corresponding to the canonical GU dinucleotide core^19^ of the cryptic *LMNA* exon 11 donor. The c.1824C>T substitution lies at +6 within the same donor motif and forms part of the guide-target sequence, creating a single-nucleotide mismatch with the wild-type allele that favours recognition of the disease allele. Indels that disrupt the GT donor core are expected to abolish pathogenic donor function without requiring precise correction of the activating variant. The mismatch with the wild-type sequence was designed to preserve canonical lamin A and lamin C expression.

In HGPS patient-derived cells, PSS reduced use of the progerin-producing cryptic splice junction, suppressed progerin production and retained lamin A and lamin C expression. A single neonatal administration of adeno-associated virus serotype 9 (AAV9) carrying the mouse guide to Cas9-expressing *Lmna*^G609G/G609G^ mice more than doubled median lifespan, attenuated cardiovascular and multiorgan pathology and enabled multiple offspring production from treated homozygous pairs.

### Cryptic splice-site disruption suppresses progerin

PSS targets the donor required for progerin-producing splicing rather than the nucleotide that activates it. Major-spliceosome 5′ splice sites comprise a short −3 to +6 motif recognised by U1 small nuclear ribonucleoprotein (snRNP), in which a +1/+2 GU dinucleotide present in approximately 99% of human donors forms the conserved donor core, while surrounding nucleotides modulate donor strength^19–21^. In the *LMNA* exon 11 cryptic donor, c.1824C>T lies at +6, increasing complementarity to U1 small nuclear RNA (snRNA) and promoting use of the pre-existing site^3,12^. We exploited this local sequence arrangement to design the human PSS single-guide RNA (hPSS-sgRNA) to pair across c.1824T while positioning Cas9 cleavage at the corresponding +1/+2 GT dinucleotide. Cleavage-associated indels were intended to abolish donor recognition. The wild-type sequence introduces a single guide–target mismatch, a configuration designed to favour editing of the mutant allele and thereby preserve canonical lamin A and lamin C expression (**Fig. 1a**).

**Fig. 1.**
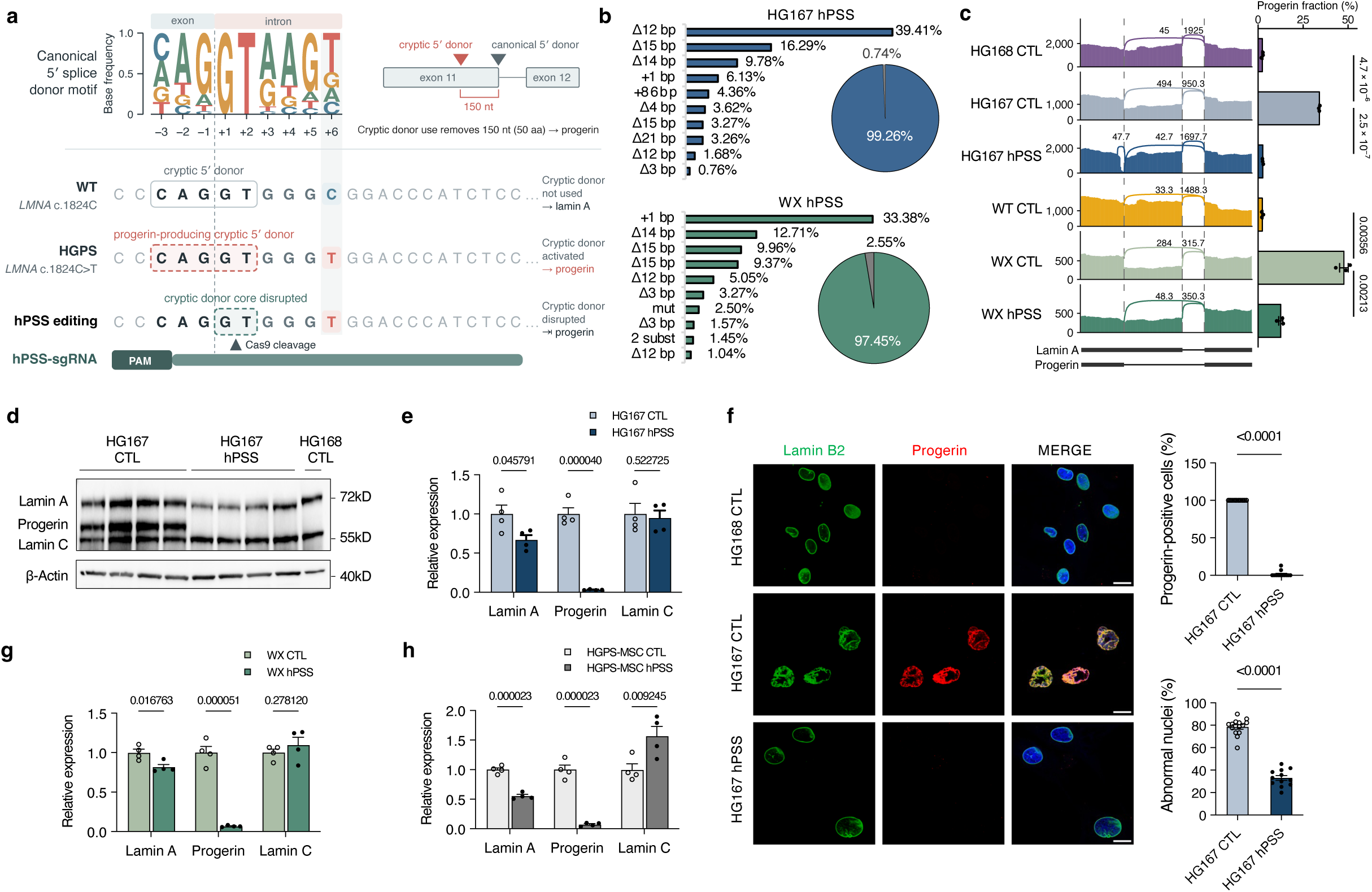
Disruption of the *LMNA* cryptic donor suppresses progerin-producing splicing in human HGPS cells. **a,** Schematic of CRISPR–Cas9 targeting of the progerin cryptic splice donor in *LMNA* exon 11. The 5′ splice-donor motif spans positions −3 to +6, with a conserved GT dinucleotide at +1/+2. The c.1824C>T mutation at +6 enhances cryptic donor use. The hPSS single-guide RNA (sgRNA) matches the mutant T, introducing one mismatch with wild-type LMNA, and positions Cas9 cleavage at the GT core to disrupt the donor through non-homologous end joining (NHEJ)-mediated insertions and deletions. **b,** Target-site sequence profiles in hPSS-treated HG167 and WX HGPS fibroblasts. Bars show the ten most abundant sequence classes among reads assigned to the HGPS-mutant allele after wild-type (WT)-sequence filtering; pies show modified and unmodified fractions. **c,** *LMNA* junction usage and progerin-junction fractions, calculated as cryptic/(cryptic + canonical) junction reads (*n* = 3 biological replicates per group). Healthy-reference HG168 and WT cells received non-targeting CTL-sgRNA (CTL). Comparisons used Welch’s t-tests with Benjamini–Hochberg correction across six comparisons; adjusted *P* values are shown. **d,e,** Representative HG167 immunoblots (d) and lamin A, progerin and lamin C abundance in HG167 fibroblasts (e). **f,** Lamin B2 (green), progerin (red) and DAPI (blue) staining in CTL- or hPSS-treated HG167 cells, with CTL-treated HG168 as a healthy reference. Progerin-positive and abnormal nuclear fractions: *n* = 14 CTL and 12 hPSS fields from three independent cultures; Student’s and Welch’s t-tests, respectively. Scale bar, 10 μm. **g,h,** Lamin A, progerin and lamin C abundance in WX fibroblasts (g) and HGPS-derived mesenchymal stem cells (h). Protein abundance in e,g,h was normalized to β-Actin and matched CTL (*n* = 4 independent cultures per group); unpaired Student’s t-tests with two-stage Benjamini–Krieger–Yekutieli correction across three proteins (target FDR, 1%); *q* values are shown. Data are mean ± s.e.m.

To test whether donor-directed editing altered the splice decision that generates progerin, we introduced Cas9 with hPSS-sgRNA or a non-targeting control guide (CTL) into HGADFN167 (HG167) and independent patient (WX) fibroblasts. Amplicon sequencing revealed heterogeneous insertions and deletions centred on the cryptic donor, with distinct predominant outcomes in the two backgrounds, a 12-bp deletion accounted for 39.4% of mutant-allele reads in HG167, whereas a 1-bp insertion accounted for 33.4% in WX (**Fig. 1b and Supplementary Table 1**). Despite these different repair spectra, both converged on reduced use of the progerin-producing junction. The progerin-junction fraction decreased from 34.0% to 2.6% in HG167, approaching the 2.3% observed in both CTL-treated HGADFN168 (HG168) and unrelated wild-type (WT) fibroblasts, and from 47.7% to 12.6% in WX fibroblasts (**Fig. 1c**). Thus, hPSS treatment introduced highly efficient donor-core-disrupting edits and reduced use of the progerin-producing splice junction.

We next quantified the lamin isoforms to determine whether the splice response reduced progerin while retaining canonical protein output. In HG167 fibroblasts, progerin decreased by 96.8%, whereas lamin A and lamin C remained at 67.0% and 95.1% of their respective control abundance (**Fig. 1d, e**). A second guide offset by one nucleotide also reduced progerin, with the stronger suppression obtained using hPSS-sgRNA supporting its selection for subsequent analyses (**Extended Data Fig. 1a, b**). Lamin B1, a nuclear-lamina component whose loss is associated with cellular senescence^22,23^, increased after treatment (**Extended Data Fig. 1c**). To examine the effect of the same guide in a healthy background, we treated HG168 fibroblasts with hPSS-sgRNA and found no detectable changes in lamin A, lamin C or lamin B1 abundance (**Extended Data Fig. 1d**).

The reduction in progerin prompted us to examine whether treatment also improved the nuclear and senescence-associated abnormalities of HGPS cells. In HG167 fibroblasts, hPSS-sgRNA reduced the progerin-positive cell fraction by 98.4% and lowered the fraction of cells with compromised nuclear integrity by 57.9% (**Fig. 1f**), together with increased cell proliferation and a lower senescence-associated β-galactosidase-positive fraction (**Extended Data Fig. 1e)**. Because loss of heterochromatin and disruption of nuclear-lamina organization are characteristic features of cellular senescence and HGPS, we also examined corresponding nuclear markers. hPSS increased heterochromatin protein 1α (HP1α), lamina-associated polypeptide 2 (LAP2), and histone H3 lysine 9 trimethylation (H3K9me3) signals, consistent with improved heterochromatin maintenance and lamina organization (**Extended Data Fig. 1f**). The cellular response therefore extended beyond progerin suppression to nuclear organization and senescence-associated phenotypes.

We next assessed whether the hPSS response was reproducible across an independent patient background and a disease-relevant mesenchymal context, as mesenchymal lineages are prominently affected in HGPS. In WX fibroblasts and HGPS patient induced pluripotent stem cell (iPSC)-derived mesenchymal stem cells (HGPS-MSCs), hPSS reduced progerin by 93.1% and 92.7%, respectively, while retaining lamin A and lamin C expression (**Fig. 1g, h and Extended Data Fig. 2a, b**). Both models reproduced improvements in nuclear integrity, senescence-associated phenotypes as well as lamina and heterochromatin organization (**Extended Data Fig. 2c–g**).

We also characterised the genome-wide sequence outcomes and cleavage specificity of hPSS. GUIDE-seq2 identified six three-mismatch candidate sites in HG167 and nine in WX, all intronic or intergenic, and no non-*LMNA* candidates with zero to two guide mismatches (**Extended Data Fig. 3a, b and Supplementary Table 2**). Whole-genome sequencing (WGS) provided complementary evidence of cut-proximal indels at a subset of these candidate sites (**Extended Data Fig. 3c, d and Supplementary Tables 3 and 4**). No coding-exon off-target editing was detected.

Together, donor-core editing redirected *LMNA* splicing away from the progerin-producing junction with high efficiency and specificity. Across two independent patient-derived fibroblast lines and iPSC-derived stem cells, hPSS reduced progerin by 93–97% while retaining canonical lamin expression and reproducibly improving multiple HGPS-associated cellular phenotypes.

### hPSS counteracts transcriptional and chromatin dysregulation

The cellular response to hPSS was underpinned by changes in disease-associated transcriptional states. In both HG167 and WX fibroblasts, hPSS opposed disease-associated expression changes and shifted gene expression towards healthy-reference profiles (**Extended Data Fig. 4a–c and Supplementary Table 5**). Within the 414 genes rescued in both HGPS fibroblast lines, hPSS increased expression of the cell-cycle-related genes *E2F2* and *MKI67* and the nuclear-lamina gene *LMNB1*, while reducing the stress-inflammatory and matrix-remodelling genes *PTGS2* and *INHBA* (**Fig. 2a and Supplementary Table 6**). These changes converged on increased chromosome-segregation, DNA-replication, E2F/G2M, and DNA-repair programs and decreased senescence-associated secretory phenotype (SASP), TNF–NF-κB, hypoxia, IFN-γ, and cytokine-response programs (**Fig. 2b and Supplementary Table 7**). hPSS therefore rescued two independent HGPS fibroblast lines toward a more proliferative and genome-maintenance-competent transcriptional state with lower senescence-, inflammatory- and stress-associated activity.

**Fig. 2.**
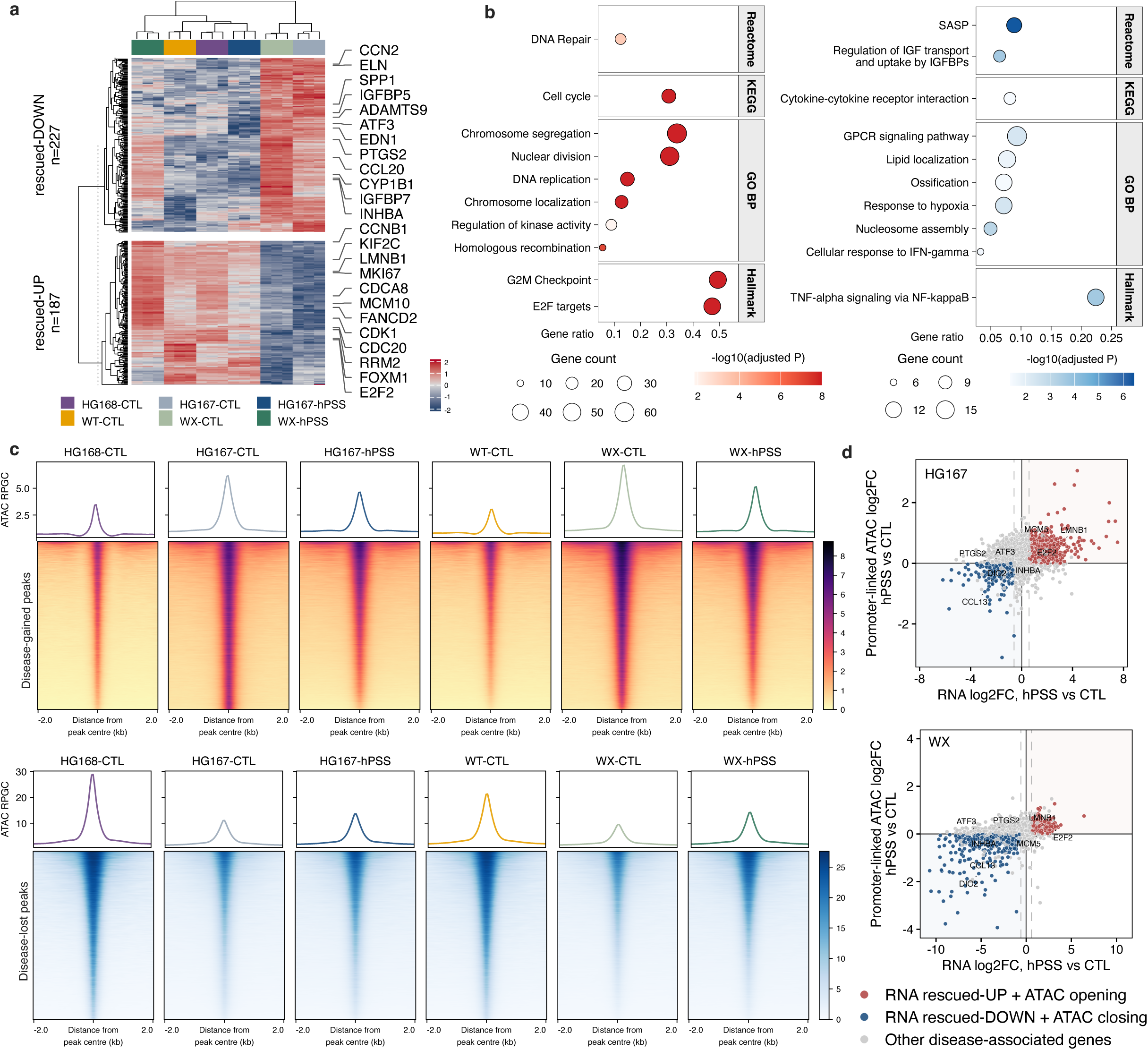
hPSS shifts disease-associated transcriptional and chromatin states towards healthy references. **a,** Gene-wise Z scores of trimmed mean of M values (TMM)-normalized log₂ counts per million for 414 genes rescued in both HG167 and WX fibroblasts (187 rescued-UP; 227 rescued-DOWN). Rescued-UP genes were suppressed relative to both non-targeting CTL-sgRNA-treated healthy references, WT and HG168, and increased with human progerin splice-site silencing sgRNA (hPSS); rescued-DOWN genes met the opposite criteria. Each comparison required adjusted *P* < 0.05 and ≥1.5-fold change. **b,** Gene Ontology Biological Process, KEGG, Reactome and MSigDB Hallmark enrichment of shared rescued-UP (left) and rescued-DOWN (right) genes. **c,** ATAC-seq profiles and heatmaps across ±2 kb of disease-gained (upper; 69,149) and disease-lost (lower; 52,044) peak centres. Sets combine peaks increased or decreased relative to both healthy references in either HGPS line. Signals are normalized to reads per genomic content (RPGC). **d,** RNA–promoter accessibility concordance. Axes show hPSS-versus-CTL log₂ fold changes; promoter accessibility is the median across linked peaks. *n* = 3 biological replicates per group for RNA-seq and ATAC-seq. Differential and enrichment analyses used DESeq2 Wald and hypergeometric tests, respectively, with Benjamini–Hochberg correction.

Given that epigenetic dysregulation is a hallmark of ageing and a prominent feature of HGPS^24,25^, we next asked whether the transcriptional recovery induced by hPSS was accompanied by correction of disease-associated chromatin accessibility. Assay for transposase-accessible chromatin with sequencing (ATAC-seq) showed that treatment reduced excess accessibility at disease-gained regions and increased accessibility at sites suppressed in HGPS fibroblasts (**Fig. 2c, Extended Data Fig. 4d–f and Supplementary Table 8**). The functional annotations distinguished two components of this response. Regions regaining accessibility lay near genes enriched for adhesion, cytoskeletal organisation, migration and Wnt-related pathways (**Supplementary Table 9**). Regions with attenuated disease-elevated accessibility shared by both patient backgrounds were enriched for AP-1/basic leucine zipper (bZIP) and TEAD-family motifs, implicating stress-responsive regulatory sequences in the chromatin response to hPSS (**Extended Data Fig. 4g and Supplementary Table 10**).

Integrated RNA-seq and ATAC-seq analysis identified a subset of disease-associated genes in which expression and promoter accessibility shifted in the same direction after hPSS treatment. Promoter-linked concordance was observed for 743 of 1,988 genes in HG167 (37.4%) and 831 of 3,478 genes in WX (23.9%; **Fig. 2d and Supplementary Table 11**). Representative loci, including *LMNB1* and *INHBA*, illustrated coordinated changes in RNA abundance and promoter-proximal accessibility (**Extended Data Fig. 4h**).

### mPSS extends lifespan

The near-complete removal of progerin and improvement of disease-associated defects across patient-derived HGPS cell models prompted us to test whether PSS could modify disease progression *in vivo* using the widely used *Lmna*^G609G/G609G^ progeroid mouse model, which recapitulates major features of human HGPS^14,26^. Although the local human and mouse sequences differ at the corresponding locus (*Lmna* c.1827C>T), progerin production in both species depends on the exon 11 cryptic 5′ donor^14,26^. We therefore adapted the guide to the homologous mouse cryptic donor (mPSS) and confirmed progerin suppression in both *Lmna*^G609G/+^ and *Lmna*^G609G/G609G^ mouse embryonic fibroblasts (**Extended Data Fig. 5a, b**). These data established mPSS activity at the homologous mouse donor and provided the basis for *in vivo* evaluation in the homozygous disease model. Cas9-expressing *Lmna*^G609G/G609G^ littermates received a single retro-orbital injection of AAV9 carrying mPSS-sgRNA at postnatal day 3 (P3; 1.5 × 10¹¹ viral genomes per mouse), with PBS-injected mice serving as controls (CTL; **Fig. 3a**). Median survival increased from 115 to 279 days, a 2.43-fold extension (CTL, n = 33; mPSS, n = 52; *P* = 6.10 × 10⁻^17^, log-rank test; **Fig. 3b**). At the follow-up cutoff, the longest observed follow-up in the mPSS-treated cohort reached 421 days.

**Fig. 3.**
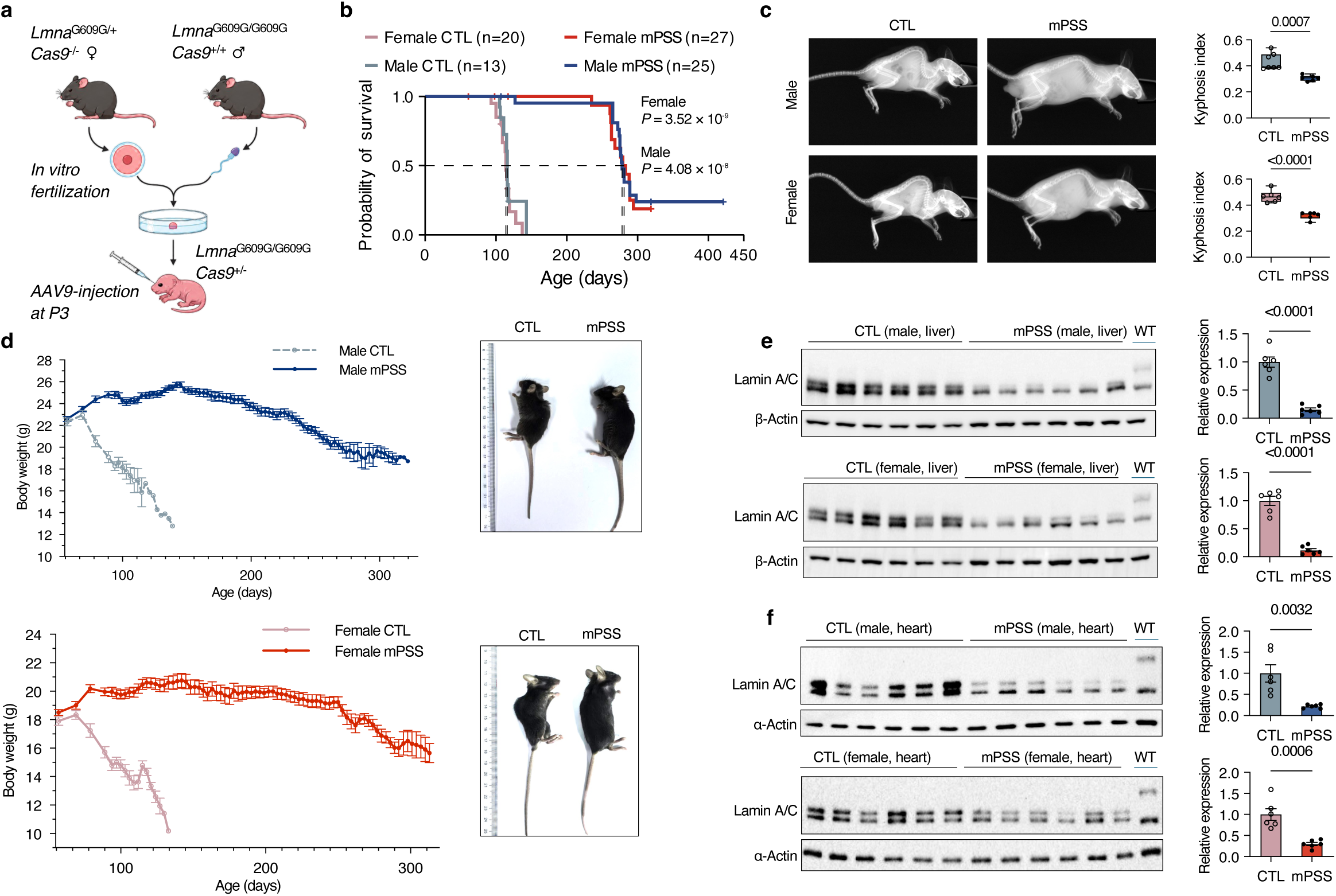
mPSS suppresses tissue progerin and prolongs survival in homozygous progeroid mice. **a,** Generation of Cas9-expressing *Lmna*^G609G/G609G^ mice by in vitro fertilization and treatment of P3 pups with AAV9 carrying mPSS-sgRNA or an equivalent volume of PBS (CTL). **b,** Sex-stratified Kaplan–Meier curves (female CTL/mPSS, *n* = 20/27; male, *n* = 13/25 mice). Ticks indicate censoring; dashed lines indicate 50% survival and median survival times. Comparisons used log-rank (Mantel–Cox) tests. **c,** Representative radiographs and kyphosis index (male CTL/mPSS, *n* = 7/6; female, *n* = 6/6 mice). Boxes show medians and interquartile ranges, with minimum-to-maximum whiskers. **d,** Longitudinal body weights and representative mouse images. Initial cohort sizes were 13 male CTL, 25 male mPSS, 20 female CTL and 27 female mPSS mice. **e, f,** Lamin A/C and progerin immunoblots and progerin abundance in liver (e) and heart (f), normalized to β-Actin and α-Actin, respectively, and expressed relative to sex-matched CTL (n = 6 tissue samples per sex, tissue and treatment). Body-weight curves and protein-abundance bars show mean ± s.e.m. Comparisons in c, e, f used unpaired Student’s *t*-tests.

mPSS also mitigated postural deformity and progressive wasting. Whole-body radiographs showed reduced kyphosis, with an approximately 30% lower kyphosis index in both sexes (**Fig. 3c**). Treated mice had greater projected body size and a higher radiograph-derived lean-area fraction (**Extended Data Fig. 5c**). Longitudinal measurements showed treated mice maintained higher body weights as CTL mice developed progressive wasting (**Fig. 3d**), and several organs were larger or heavier after treatment (**Extended Data Fig. 5d**). Random-fed blood glucose also increased in both sexes (**Extended Data Fig. 5e**). These findings show that mPSS prolonged lifespan while preserving whole-body condition over time.

At the molecular level, mPSS suppressed progerin across multiple affected tissues. Progerin abundance decreased by 84.6–88.2% in liver and 70.3–77.8% in heart across both sexes (**Fig. 3e, f**). Kidney progerin was also reduced, whereas lung levels changed little (**Extended Data Fig. 5f**), consistent with tissue-dependent editing efficiency following systemic AAV9 delivery^16,18^.

### mPSS preserves aortic structure and reduces multiorgan pathology

The systemic benefit extended to the aorta, a critical disease target because vascular smooth-muscle-cell (VSMC) loss and progressive arterial-wall pathology are major features of HGPS and contribute directly to the cardiovascular complications that drive mortality^4,27^. We examined VSMC loss and adventitial expansion to determine whether treatment preserved the vessel-wall compartments affected by the disease. Aortic VSMC density was 1.85-fold and 1.42-fold that of controls in treated males and females, respectively, while adventitial area was reduced by 71.5% and 69.0%, with better-preserved vessel-wall architecture (**Fig. 4a**). mPSS therefore attenuated both medial cell loss and adventitial remodelling.

**Fig. 4.**
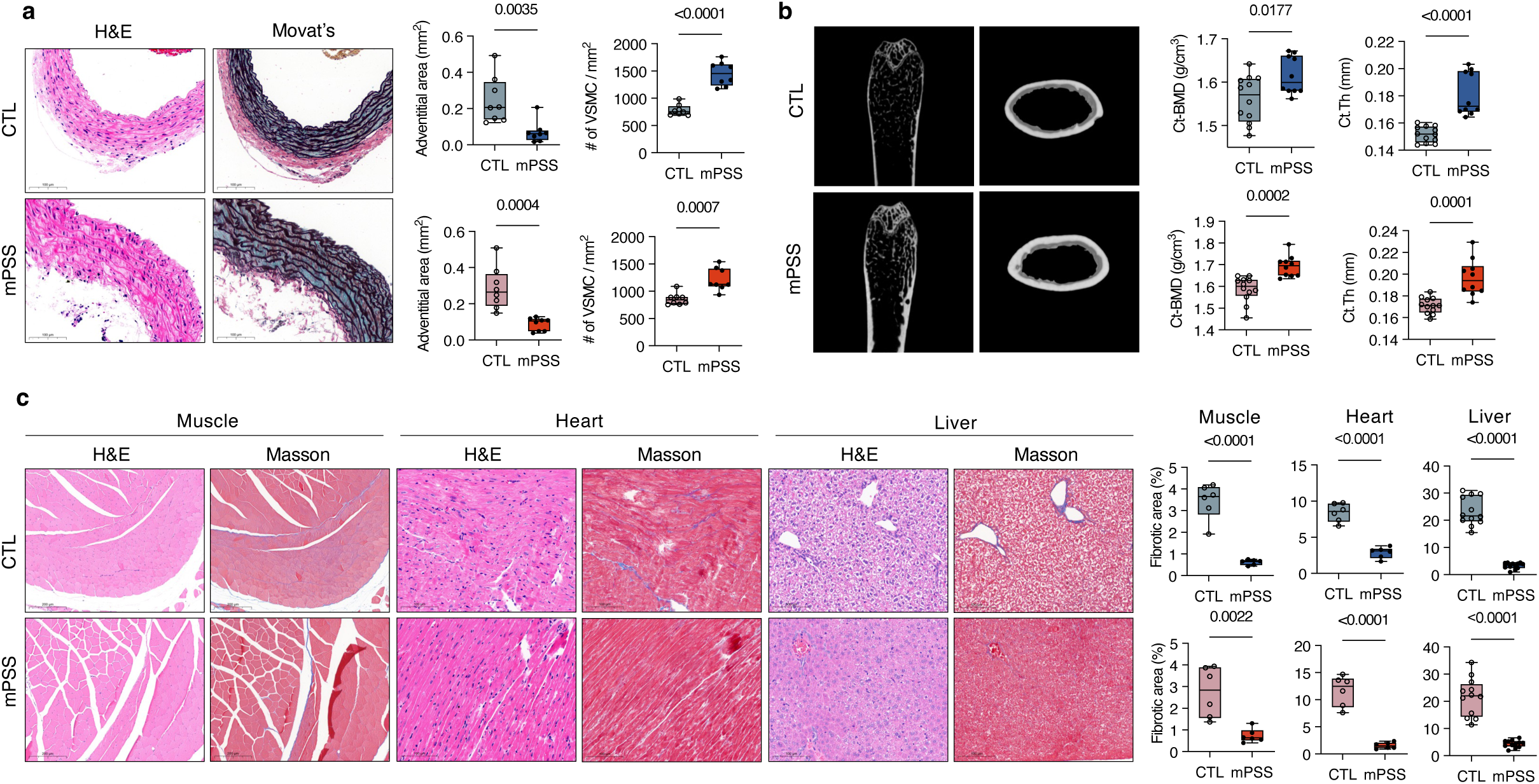
mPSS improves vascular and skeletal phenotypes and reduces tissue fibrosis. **a,** Aortic H&E and Movat’s pentachrome staining, with adventitial area and vascular smooth muscle cell (VSMC) density in *Lmna*^G609G/G609G^ mice receiving PBS (CTL) or mPSS (*n* = 8 samples per sex and treatment). Scale bars, 100 μm. **b,** Femoral microcomputed tomography images, cortical bone mineral density (Ct-BMD) and cortical thickness (Ct.Th) (CTL, n = 12 femora; mPSS, n = 10 femora per sex). **c,** H&E and Masson’s trichrome staining of skeletal muscle, heart and liver, with fibrotic-area quantification. Per sex and treatment, *n* = 6 tissue samples for muscle and heart and 12 for liver. Scale bars: muscle, 200 μm; heart, 100 μm; liver 100 μm. Quantification shows males (upper) and females (lower) in a–c. Boxes indicate medians and interquartile ranges, with minimum-to-maximum whiskers. Comparisons used unpaired Student’s *t*-tests.

We next examined bone microarchitecture to resolve the skeletal effects beyond the postural improvement. Femoral microcomputed tomography showed increased cortical bone mineral density and cortical thickness in both sexes (**Fig. 4b**). Cortical area and cortical-area fraction also increased, marrow-cavity area decreased and growth plates were thicker in treated mice (**Extended Data Fig. 6a, b**).

Treatment also reduced fibrotic pathology across multiple tissues. Fibrotic area decreased in both sexes, by 73.1–82.3% in skeletal muscle, 65.8–86.6% in heart and 79.7–86.0% in liver (**Fig. 4c**). Skin sections showed a lower collagen volume fraction and greater epidermal thickness (**Extended Data Fig. 6c**). Together, these findings demonstrate that mPSS preserved aortic and cortical-bone structure and reduced fibrosis across muscle, heart, liver and skin, providing tissue-level evidence of systemic disease modification.

### mPSS attenuates cardiac and hepatic stromal remodelling

The reduction in cardiac and hepatic fibrosis after mPSS treatment (**Fig. 4c**) led us to examine the corresponding cellular and transcriptional changes in heart and liver by single-cell RNA sequencing. In the heart, fibroblasts accounted for a smaller fraction of recovered cells after treatment, decreasing from 62.1% to 39.3%, whereas the endothelial fraction increased from 25.2% to 44.5% (**Fig. 5a, b, Extended Data Fig. 7a–e and Supplementary Table 12**). Transcriptional analysis showed reduced inflammatory and vascular-remodelling programmes across several cardiac populations, including lower TNF–NF-κB and TGF-β signalling and reduced stress-response programmes (**Fig. 5c, Extended Data Fig. 7f and Supplementary Tables 13 and 14**).

**Fig. 5.**
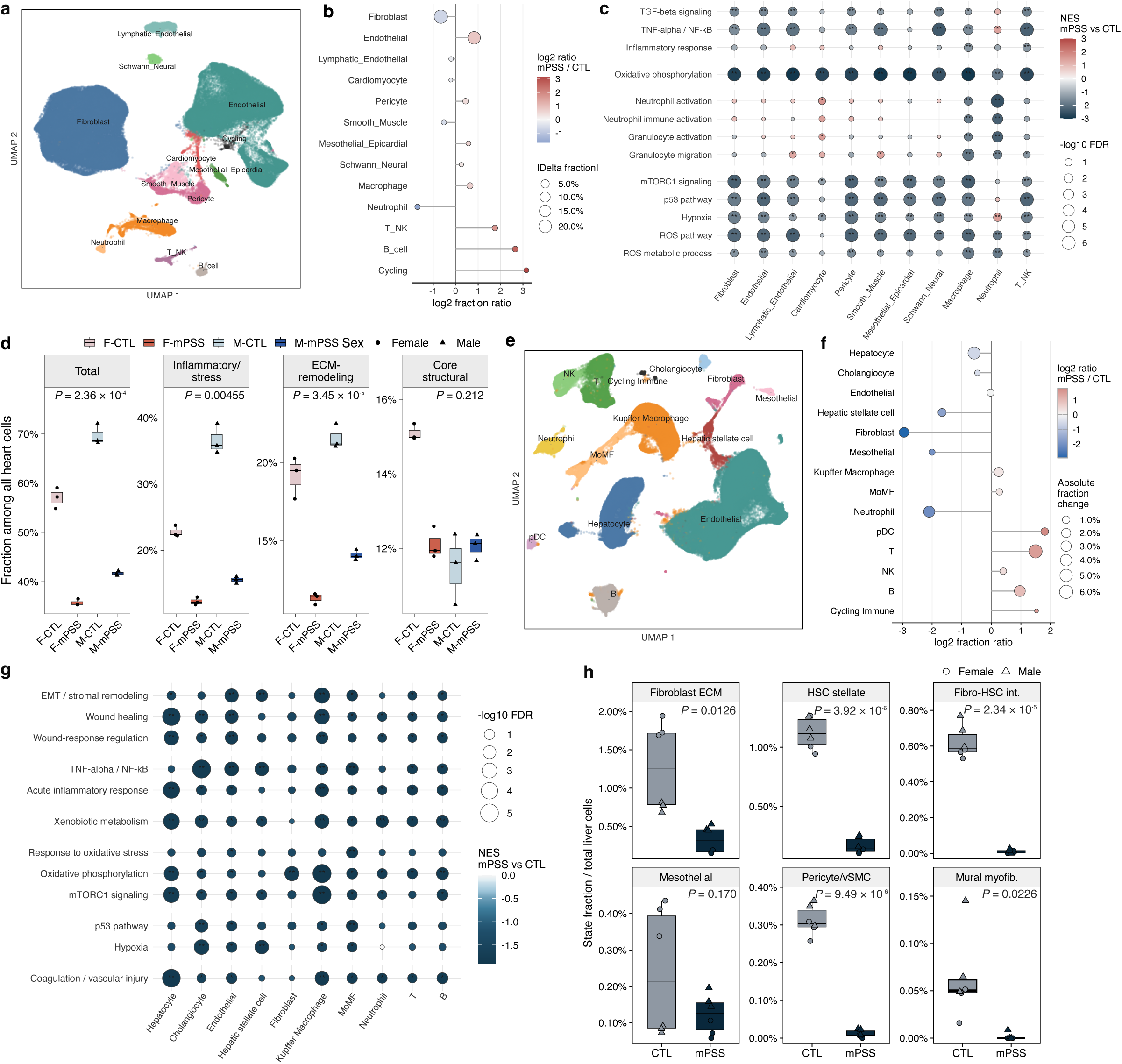
mPSS reduces cardiac and hepatic remodelling-associated cell states. **a,** UMAP of 114,186 recovered heart cells from CTL- and mPSS-treated *Lmna*^G609G/G609G^ mice. **b,** Cardiac cell-type fractions pooled across libraries within each treatment. **c,** Cardiac gene-set enrichment. **d,** Total fibroblast and fibroblast-state fractions among all recovered heart cells. **e–g,** UMAP (**e**), cell-type fractions (**f**) and gene-set enrichment (**g**) of 157,979 recovered liver cells from CTL- and mPSS-treated *Lmna*^G609G/G609G^ mice. MoMF, monocyte-derived macrophages; pDC, plasmacytoid dendritic cells; EMT, epithelial–mesenchymal transition. **h,** Hepatic stromal-state fractions among all recovered liver cells. HSC, hepatic stellate cell; vSMC, vascular smooth muscle cell; int., intermediate; myofib., myofibroblast. In **c,g**, preranked gene-set enrichment used Benjamini–Hochberg correction; positive normalized enrichment scores are displayed at zero in **g**. *, 0.05 ≤ FDR < 0.25; **, FDR < 0.05. In **d,h**, points denote libraries (n = 6 per treatment); boxes show medians and interquartile ranges (IQRs), with whiskers extending to values within 1.5 IQRs. Treatments were compared with sexes pooled using Welch’s *t*-tests with Benjamini–Hochberg correction across four measures in **d** and six states in **h**; adjusted *P* values are shown.

We then examined which stromal states accounted for these compositional changes. Inflammatory/stress and extracellular matrix (ECM)-remodelling fibroblasts decreased from 29.7% to 13.9% and from 20.4% to 12.7% of recovered cells, respectively. Both reductions were evident in males and females, whereas the core structural fraction showed no detectable difference (**Fig. 5d and Extended Data Fig. 7g, h**). An independent, cluster-free MiloR^28^ analysis localized CTL-enriched neighborhoods to the inflammatory/stress state, further supporting this change (**Extended Data Fig. 7i and Supplementary Table 15**).

In the liver, hepatic stellate cells, fibroblasts and neutrophils accounted for smaller fractions of recovered cells after treatment (**Fig. 5e, f and Extended Data Fig. 8a–e**). Reduced wound-response, fibrogenic and inflammatory expression programmes further characterised the hepatic response (**Fig. 5g and Extended Data Fig. 8f**). Within the stromal compartment, ECM-remodelling fibroblast and stellate-like fractions decreased, while fibroblast–stellate intermediate, contractile mural and myofibroblast-like states became rare after treatment (**Fig. 5h, Extended Data Fig. 8g–i and Supplementary Table 16**). Retained stromal populations showed lower inflammatory and niche-associated gene expression, while structural ECM genes showed mixed responses (**Extended Data Fig. 8j, k**). CellChat^29^ inferred reduced inflammatory communication towards cardiac fibroblasts and hepatic stromal cells, and liver myeloid cells showed lower *Cxcl2* and *Mif* expression (**Extended Data Figs. 7j, k and 8l–n**). Together, these cell-level analyses support the histological reduction in cardiac and hepatic fibrosis by identifying smaller fractions of remodelling-associated stromal states and lower inflammatory and stress-related transcription after mPSS treatment.

### mPSS enables offspring production from *Lmna*^G609G/G609G^ pairs

Previous studies reported severe reproductive impairment in *Lmna*^G609G/G609G^ mice, and successful offspring production from *Lmna*^G609G/G609G^ pairings has not been reported^14,26^. These infertility records made successful breeding by treated homozygous pairs a stringent functional test of recovery. Every mPSS-treated *Lmna*^G609G/G609G^ breeding pair produced offspring, yielding 52 pups across seven litters. Deliveries occurred between P82 and P137, with three litters born after the 115-day median lifespan of CTL mice (**Fig. 6a and Supplementary Table 17**). Of the 52 offspring, 37 were delivered vaginally and 15 by Caesarean section after dystocia, and newborn pups were transferred to lactating ICR foster dams (**Fig. 6b**). To our knowledge, this is the first reported offspring production from pairings in which both parents were homozygous for *Lmna*^G609G/G609G^ following a postnatal intervention.

**Fig. 6.**
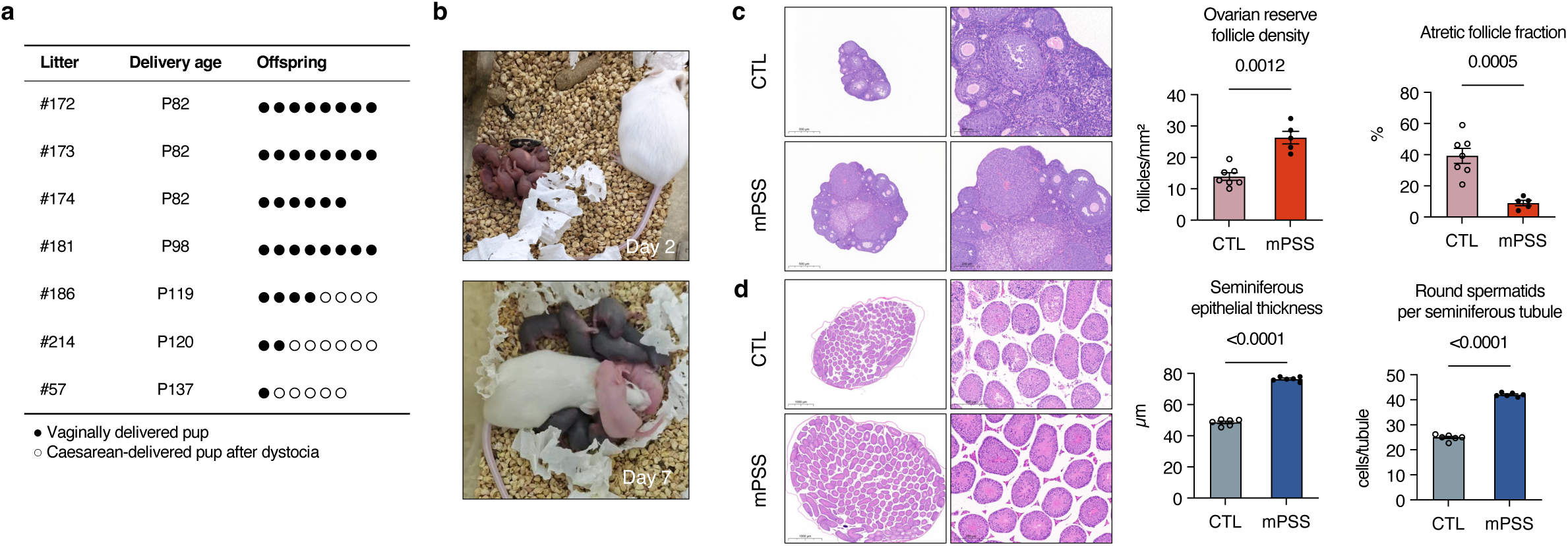
mPSS enables offspring production and improves reproductive tissues. **a,** Breeding outcomes of *Lmna*^G609G/G609G^ mice treated with mPSS. Delivery age denotes maternal postnatal (P) age. **b,** Offspring at postnatal days 2 and 7 with lactating ICR foster dams. **c,** Ovarian H&E sections, ovarian reserve-follicle density and atretic-follicle fraction in PBS-treated controls (CTL) and mPSS-treated mice (CTL, *n* = 7; mPSS, *n* = 5 ovarian samples). Scale bars are indicated. **d,** Testicular H&E sections, seminiferous epithelial thickness and round spermatids per seminiferous tubule. Three fields were averaged per mouse (*n* = 6 mice per group). Scale bars: overview, 1,000 μm; enlarged views, 200 μm. Data are mean ± s.e.m. Comparisons used Welch’s *t*-tests.

PCR genotyping confirmed the *Lmna*^G609G/G609G^ genotype in all 52 offspring, and target-region Sanger sequencing detected no edited allele in the eight offspring examined from one litter (**Extended Data Fig. 9a–c**). Twenty offspring survived for long-term follow-up and developed the expected progeroid phenotype, with a median survival of 114 days (**Extended Data Fig. 9d**).

The reproductive outcome was supported by improvement in both female and male gonads. Gross measurements showed larger ovaries and increased testis weight after mPSS treatment (**Extended Data Fig. 9e, f**). In the ovary, mPSS preserved the follicular reserve, with higher reserve-follicle density and lower follicular atresia, together with supporting improvements in follicle composition and stromal fibrosis (**Fig. 6c and Extended Data Fig. 9g**). In the testis, greater seminiferous epithelial thickness and increased round spermatids indicated improved spermatogenic progression, with additional histomorphometry supporting better seminiferous organization (**Fig. 6d and Extended Data Fig. 9g, h**).

Repeated offspring production, together with improvement of testicular and ovarian architecture, demonstrates recovery of reproductive capacity after mPSS treatment.

## Discussion

PSS establishes the progerin-producing cryptic donor as a genomic target for sustained disease modification in progeria. In human HGPS cells, donor-directed editing suppressed pathogenic splicing and progerin production while retaining lamin A and lamin C expression. Despite differences in the local human and mouse target sequences, targeting the corresponding pathogenic donor reproduced the therapeutic principle established in human cells. A single postnatal intervention at the homologous mouse cryptic donor prolonged median lifespan from 115 to 279 days and improved vascular, somatic and reproductive outcomes. These findings provide in vivo evidence that DNA-level disruption of a pathogenic splice element can alter the systemic course of a lethal progeroid disease. The activating mutation need not be precisely corrected when the splice signal responsible for its pathogenic output can instead be disabled.

PSS converts splice suppression into a durable genomic intervention while retaining expression of both canonical *LMNA* isoforms. Earlier CRISPR–Cas9-based disruption reduced progerin by disrupting the shared lamin A/progerin coding region^16,17^. Donor targeting instead directs disruption towards pathogenic splice-site function. Although lamin A depletion is tolerated in mouse models^30^, its dispensability across human tissues remains unresolved. Evidence for non-redundant functions of lamin A and lamin C provides a biological rationale for retaining both during sustained progerin suppression. Lamin A contains a nucleosome-binding motif absent from lamin C^31^ and supports SIRT6-dependent DNA repair^32^, whereas lamin C has a specific role in re-establishing genome organisation after mitosis^33^. The continued expression of both isoforms in our human models therefore represents a biologically significant feature of donor-directed treatment.

PSS also differs from mutation correction in what constitutes a productive edit. Correction of c.1824C>T requires a defined nucleotide conversion, whereas donor disruption can use different repair sequences that disable pathogenic splice-site function. Cas9-induced end joining generates heterogeneous insertions and deletions, converging on the same functional endpoint of eliminating pathogenic splice-site activity (**Fig. 1b, c**). PSS therefore combines sustained genome modification with a broader productive repair space centred on loss of donor function.

The therapeutic value of this donor-directed design was evident in the magnitude and breadth of the in vivo response. In Cas9-expressing *Lmna*^G609G/G609G^ mice, a single neonatal administration of AAV9-mPSS increased median lifespan from 115 to 279 days, a 143% increase. The closest guide-delivery precedent reported a 26.5% increase with constitutive Cas9 expression and neonatal AAV9 administration^16^, while delivery of Cas9 and guide RNA together produced a 26.4% increase^17^. In a distinct human-*LMNA* transgenic model, adenine base editing increased median survival from 189 to 337 days after P3 administration, a 1.78-fold increase compared with the 2.43-fold increase observed here^18^. Beyond longevity, mPSS preserved aortic wall structure, improved cortical bone architecture and reduced fibrosis across multiple tissues. Treated pairs in which both parents were homozygous produced 52 offspring across seven litters, establishing a reproductive outcome not reported in these earlier editing studies.

The present study establishes systemic efficacy of PSS through neonatal AAV9 delivery of mPSS-sgRNA to Cas9-expressing mice. Clinical application will require delivery of both the nuclease and guide. Compact Cas9 systems such as Staphylococcus aureus Cas9 (SaCas9) and Campylobacter jejuni Cas9 (CjCas9) have already enabled therapeutic genome editing in vivo^34^, including in HGPS models^17^, providing a basis for adapting PSS to a self-contained vector. The vascular and multiorgan responses define disease-relevant endpoints for refining tissue delivery and assessing efficacy across doses and treatment ages, including after disease onset. Guide optimisation and high-fidelity Cas9 variants offer routes to reduce unintended editing at the candidate loci identified by GUIDE-seq2 and WGS^35^. Together, the activity in patient-derived cells and sustained in vivo benefits support clinical development of a one-time donor-directed treatment that suppresses progerin while retaining canonical lamin expression.

More broadly, PSS defines pathogenic splice signals as targets for durable genomic intervention. The corresponding human and mouse donors required different guide sequences, but the same functional targeting principle suppressed progerin in both settings. This approach warrants investigation in other disorders where a discrete pathogenic splice signal can be disrupted while preserving essential transcript and protein outputs. In cystic fibrosis, the deep-intronic *CFTR* c.3718-2477C>T variant creates a cryptic donor that promotes pseudoexon inclusion and reduces productive *CFTR* expression^7^; in collagen VI-related dystrophy, *COL6A1* c.930+189C>T activates a cryptic donor that drives inclusion of a dominant-negative pseudoexon^36^. RNA-level blockade of these events restores *CFTR* channel activity and collagen VI matrix formation, respectively, demonstrating that selective suppression of the aberrant splice product can recover disease-relevant function^37,38^. The principle may also extend beyond mutation-defined inherited lesions. In castration-resistant prostate cancer, cryptic-exon inclusion generates the constitutively active androgen receptor splice variant 7 (AR-V7) isoform, which is associated with resistance to androgen-receptor pathway inhibition; disruption of the cis-elements required for AR-V7 production could therefore provide a route to isoform-selective suppression^39^. Together, these examples point to a broader therapeutic opportunity in treating pathogenic splice signals as targetable cis-regulatory elements, extending genome editing beyond variant correction toward durable control of disease-causing isoform production.

## Acknowledgments

This work was supported by grants of Science, Technology and Innovation Commission of Shenzhen Municipality (JCYJ20210324,114408024), and Guangdong Basic & Applied Basic Research Foundation (2021B1515130004).

## Author contributions

W.J. and J.M. conceived the project. W.J. and J.M. designed the experiments. W.J. did the experiments with help from H.Y., Y.Z., Z.X.Z., S.K.K.W, and T.L. J.M. performed the bioinformatic analysis with help from J.H.Z. H.Y. monitored and recorded animal-related work. X.L. helped with data discussion. W.J. and J.M. interpreted the data. W.J. and J.M. prepared the manuscript with comments and inputs from all authors. Z.Z. provided the funding for this project. All authors read and approved the final manuscript.

## Data and code availability

The RNA-seq, ATAC-seq, GUIDE-seq-2, and mouse scRNA-seq datasets generated in this study have been deposited in the NCBI Gene Expression Omnibus under SuperSeries accession GSE341429, comprising GSE341425, GSE341424, GSE341427, and GSE347409, respectively. WGS and amplicon sequencing data are deposited in the NCBI Sequence Read Archive under BioProject accession PRJNA1501649. All datasets will be released publicly upon publication. Data analyses used published software and packages, with analysis parameters specified in Methods.

## Methods

### Cell culture

Human primary fibroblasts HGADFN167 (HG167) and HGADFN168 (HG168) were obtained from the Progeria Research Foundation (PRF). Primary fibroblasts from a healthy adult donor (wild type, WT) and a Hutchinson–Gilford progeria syndrome (HGPS) patient (WX) were established in our laboratory and described previously^1^. Human primary fibroblasts were cultured in DMEM supplemented with 15% (v/v) Foundation™ Fetal Bovine Serum (FBS) (GeminiBio), 1% Non-Essential Amino Acids Solution, 1% GlutaMAX, 1% penicillin/streptomycin and 10 ng/ml basic FGF (PeproTech). WT and HGPS mesenchymal stem cells (MSCs) were derived from induced pluripotent stem cells and maintained in alpha-MEM-based medium as described previously^1^. HEK293T cells were cultured in antibiotic-free DMEM supplemented with 10% FBS (Gibco). Mouse embryonic fibroblasts (MEFs) were isolated from E13.5 embryos and maintained in DMEM supplemented with 10% FBS. All cells were cultured at 37 °C in 5% CO2 and tested negative for mycoplasma by PCR at the Centre for PanorOmic Sciences, Li Ka Shing Faculty of Medicine.

### Plasmids and sgRNA cloning

The single-guide RNAs (sgRNAs) were designed to target the progerin-specific splice-donor sequence adjacent to a Streptococcus pyogenes Cas9 NGG protospacer-adjacent motif (PAM): human progerin splice-site silencing (hPSS)-sgRNA (5′-GGAGATGGGTCCACCCACCT-3′), hPSS-sgRNA-2 (5′-AGGAGATGGGTCCACCCACC-3′), and mouse progerin splice-site silencing (mPSS)-sgRNA (5′-GGAGATGGATCCACCCACCT-3′). A non-targeting sgRNA (5′-CGCTGGAGAGCAACTGCATA-3′) served as the control. Oligonucleotides were synthesized by IDT, annealed, and ligated into BsmBI-digested lentiCRISPR v2 (Addgene #52961). Clones were screened and confirmed by Sanger sequencing.

### Lentivirus packaging and transduction

Lentivirus was produced by cotransfecting 15 μg lentiCRISPR plasmid, 12 μg psPAX2, and 3 μg pMD2.G (Addgene #12259 and #12260) into HEK293T cells in 15-cm dishes using 120 μg polyethyleneimine (Sigma). Supernatants collected at 48 and 72 h after transfection were filtered through 0.45-μm PVDF membranes and concentrated by ultracentrifugation at 35,000 rpm for 3 h at 4°C. Pellets were resuspended in 1 ml cold PBS, aliquoted, and stored at −80°C. For transduction, viral stocks were thawed on ice and added to culture medium containing 8 μg/ml polybrene. Medium was replaced the next day, and 2 μg/ml puromycin was added 2 days later. Cells were maintained under selection for 2 weeks before genomic DNA, RNA, and protein were collected.

### Western blotting

Cultured cells were lysed for 30 min on ice in RIPA buffer (50 mM Tris-HCl, pH 7.4; 150 mM NaCl; 1% NP-40; 1% sodium deoxycholate; 0.1% SDS; 5 mM EDTA) with protease inhibitors and clarified at 13,000 rpm for 15 min at 4°C. Frozen tissues (10–30 mg) were homogenized in RIPA buffer with a TissueLyser II and 5-mm bead for 30 s at 25 Hz, extracted on ice for 45 min, and clarified identically. Protein concentration was measured by BCA assay. Proteins were resolved on 4–12% Bis-Tris or 8% Tris gels, transferred to PVDF, blocked, and incubated with primary antibodies overnight at 4°C and HRP-conjugated secondary antibodies for 1 h at room temperature. Signals were detected with a Bio-Rad ChemiDoc and quantified in ImageJ. Primary antibodies were lamin A/C (A19524, Abclonal; 1:5,000), lamin B1 (AC057; 1:8,000), α-actin (A2235; 1:50,000), and β-actin (AC026; 1:50,000). Secondary antibodies were anti-rabbit AS014 and anti-mouse AS003 (Abclonal; 1:5,000).

### Immunostaining

Cells were seeded on 18-mm coverslips, fixed with 4% paraformaldehyde for 10 min, and blocked for 1 h in PBS containing 0.5% Triton X-100, 1% FBS, and 5% BSA. Primary antibodies were applied overnight at 4°C, followed by fluorescent secondary antibodies for 1 h at room temperature and DAPI counterstaining. Images were acquired on a Zeiss LSM 880 confocal microscope, processed in ZEN Blue, and quantified in ImageJ. Primary antibodies were lamin A/C (sc-376248, Santa Cruz; 1:150), progerin (SAB4200272, Sigma; 1:200), lamin B2 (ab151735, Abcam; 1:200), heterochromatin protein 1α (HP1α; ab109028, Abcam; 1:200), lamina-associated polypeptide 2 (LAP2; ab223768, Abcam; 1:100), and histone H3 lysine 9 trimethylation (H3K9me3; ab8898, Abcam; 1:500). Secondary antibodies were Alexa Fluor 488 goat anti-rabbit IgG and Alexa Fluor 568 goat anti-mouse IgG (Thermo Fisher Scientific; 1:1,000).

### Senescence and crystal violet staining

Senescence-associated β-galactosidase (SA-β-gal) activity was measured with a commercial kit (C0602, Beyotime) according to the manufacturer’s protocol. Cells were fixed for 5 min, incubated with staining solution overnight at 37°C without CO2, and rinsed with PBS. For crystal-violet staining, 5,000 or 10,000 cells per well were cultured in 12-well plates for 7–12 days, fixed with 4% paraformaldehyde, stained with 0.1% crystal violet for 10 min, washed, and air-dried. Cells were counted from bright-field images in ImageJ. SA-β-gal-positive fractions were calculated per field.

### CRISPR indels at the PSS site

Genomic DNA was extracted following two weeks of antibiotic selection using a commercial kit according to the manufacturer’s instructions (Qiagen). Target loci were initially amplified by PCR using PrimeSTAR GXL DNA Polymerase (Takara) with the following primers: forward, *`5′-`*TCGTCGGCAGCGTCAGATGTGTATAAGAGACAGCGTCCCGCCTGAGCCTTGTCT-3′; reverse, 5′-GTCTCGTGGGCTCGGAGATGTGTATAAGAGACAGCTGGAGTTGCCCAGGAGGTAG-3′. For high-resolution analysis, PCR amplicons were processed for deep sequencing using Nextera standard indexed adapters. Libraries were sequenced on an Illumina platform with 250-bp paired-end reads.

### RNA-seq library preparation

RNA-seq libraries were prepared using 0.5 μg of total RNA per sample with the VAHTS Universal V6 RNA-seq Library Prep Kit for Illumina (NR604, Vazyme), following the manufacturer’s instructions. Briefly, mRNA was enriched using poly-T oligo-attached magnetic beads and fragmented. First-strand cDNA was synthesized using random hexamer primers, followed by second-strand cDNA synthesis using DNA polymerase I and RNase H. The resulting double-stranded cDNA was purified using AMPure XP beads. Subsequent steps included end repair, A-tailing, adapter ligation, and size selection, followed by PCR enrichment to generate the final sequencing libraries. Unique index codes were added to each sample for downstream sequencing. Three independent biological RNA-seq libraries were generated per group.

### Genome-wide off-target profiling

Genome-wide off-target profiling followed the GUIDE-seq-2 workflow^2^, with an 8-nucleotide unique molecular identifier (UMI) added to the Tn5 transposase adapter; double-stranded oligodeoxynucleotide (dsODN) capture followed tagmentation-based tag integration site sequencing (TTISS)^3^. In brief, one million cells were transfected with PX459 carrying the hPSS-sgRNA and a dsODN tag by nucleofection. After 72 h, genomic DNA was extracted and subjected to tagmentation using Tn5 transposase pre-loaded with sequencing adapters (Tn5-UMI-F: 5′-ACTCTTTCCCTACACGACGCTCTTCCGATCTNNNNNNNNAGATGTGTATAAGAGACAG-3′, where N denotes A, T, C, or G; Tn5-Rev: `5′-[phos]CTGTCTCTTATACACATCT-3′`) and unique molecular identifiers (UMIs). This single-step reaction simultaneously fragmented DNA and added Illumina-compatible adapters. Tagmented DNA was purified with AMPure XP beads, and dsODN-containing fragments were selectively amplified by PCR using dsODN-specific primers. Libraries were size-selected to 250-500 bp and sequenced on an Illumina platform with 150-bp paired-end reads.

### Whole-genome sequencing

Genomic DNA extracted as described above was used for whole-genome sequencing library preparation. Approximately 200 ng of genomic DNA was tagmented with 2 μl of 5 μM in-house Tn5 transposase. Tagmented DNA was purified with a MinElute kit (Qiagen), indexed with Nextera adapters, and amplified for six PCR cycles with NEBNext High-Fidelity 2× PCR Master Mix (New England Biolabs). Libraries underwent double-sided size selection with AMPure XP beads, and library quality and size distribution were assessed with an Agilent 2100 Bioanalyzer. Libraries were sequenced by Novogene on an Illumina NovaSeq platform with 150-bp paired-end reads.

### ATAC-seq library preparation

Bulk assay for transposase-accessible chromatin with sequencing (ATAC-seq) libraries were prepared as described previously^4^. Approximately 50,000 viable cells were used to isolate nuclei according to the Omni-ATAC protocol. Nuclei were resuspended in 76 μl of 0.33× PBS and combined with 20 μl of 5× TAPS-DMF buffer and 4 μl of 5 μM in-house Tn5 transposase. Tagmentation was performed in a thermomixer for 30 min at 37°C and 500 rpm. Transposed DNA was purified with a MinElute kit (Qiagen), indexed with Nextera adapters, and amplified for nine PCR cycles with NEBNext High-Fidelity 2× PCR Master Mix (New England Biolabs). Libraries underwent double-sided size selection with AMPure XP beads, were assessed with an Agilent 2100 Bioanalyzer, and were sequenced by Novogene on an Illumina NovaSeq platform with 150-bp paired-end reads.

### AAV9-mPSS production and purification

Adeno-associated virus serotype 9 (AAV9)-mPSS was produced and purified by Weizhen Biosciences (Shandong, China) using a standard helper-free triple-plasmid transfection system in HEK293T cells, followed by iodixanol density-gradient ultracentrifugation. Briefly, HEK293T cells were seeded in 10-cm dishes and grown to 80–90% confluence before cotransfection with the recombinant AAV9-mPSS genome plasmid, the pAAV rep/cap packaging plasmid, and the Ad Helper plasmid. At 72 h post-transfection, the culture supernatant and cell pellet were harvested. Viral particles in the supernatant were precipitated with PEG8000, and particles retained in the cell pellet were released by lysis; the two fractions were pooled and purified by iodixanol density-gradient ultracentrifugation. The virus-containing fraction was collected and concentrated by ultrafiltration using 100-kDa molecular-weight-cutoff columns. Viral genome titers were determined by quantitative PCR (qPCR) using a SYBR Green-based detection system (Clontech AAVpro Titration Kit v.2, Takara, 6233). Purified AAV samples were diluted 10-fold, treated with DNase I (37 °C, 30 min), heat-inactivated (95 °C, 5 min), and digested with proteinase K (37 °C, 30 min) to release encapsidated genomes. After dilution, qPCR was performed using primers targeting the inverted terminal repeat (ITR), and absolute copy number was determined from a plasmid standard curve. Viral purity was assessed by silver-stained SDS-PAGE (12% polyacrylamide gel), which resolved the VP1, VP2, and VP3 capsid proteins. Only preparations with >90% purity and a titer ≥1 × 10¹³ vg/mL were used.

### Mouse strains and animal care

The *Lmna^G^*^609^*^G/G^*^609^*^G^* mice were kindly provided by Professor Baohua Liu (Shenzhen University). *Rosa26*-CAG-Cas9 mice on a C57BL/6 background were obtained from Shanghai Model Organisms Center. To generate Cas9-expressing *Lmna^G^*^609^*^G/G^*^609^*^G^* mice, sperm from *Cas9^+/+^ Lmna^G^*^609^*^G/G^*^609^*^G^* males was used to fertilize oocytes collected from *Lmna^G^*^609^*^G/+^* females in vitro. For neonatal treatment, AAV9 carrying the mPSS-sgRNA was diluted with PBS to 1 × 10^13^ vg/mL. Mice received 15 μl of vector by retro-orbital injection at postnatal day 3. Control littermates received an equivalent volume of PBS by the same route. Body weight was recorded longitudinally for each mouse. For females used for breeding, survival was censored at first mating, and body-weight measurements from that day onwards were excluded. Mice alive at scheduled sacrifice or the last follow-up were censored. Blood glucose was measured at 10:00 in the morning under non-fasted, random-fed conditions using a Verio blood glucose meter and was analysed separately for males and females. Because biological dams did not reliably rear their litters, newborn pups were transferred to lactating outbred ICR foster dams for subsequent monitoring. All animal procedures were performed in accordance with institutional guidelines and approved under 5946-21 by the Committee on the Use of Live Animals in Teaching and Research (CULATR) at The University of Hong Kong, BG-AMS-250410-CH01-Y by Bojin Biotechnology, and J2024-0011 by Shanghai Model Organisms Center. Experimental mice were maintained on a C57BL/6 background. Mice were housed under a 12-h light–12-h dark cycle with ad libitum feeding. Tissues for phenotypic analyses were collected at 15 weeks of age unless otherwise specified.

### Histology

At designated time points, skeletal muscle, heart, liver, skin, aorta, testes, and ovaries were fixed, paraffin-embedded, sectioned, and stained with hematoxylin and eosin (H&E), Masson’s trichrome, or Movat’s pentachrome. H&E staining used hematoxylin and eosin (B1003, Wuhan Bioqiandu Biotechnology); Masson’s trichrome used kit B1011; and Movat’s pentachrome used Alcian blue, elastic van Gieson, safranin, and Sirius red reagents (C1009). Sections were dehydrated, cleared, mounted, and imaged on a Nikon Eclipse Ci or E100 microscope with a Nikon DS-U3 system. Quantification was performed in ImageJ.

### X-ray and micro-CT imaging

Whole-body radiographs were acquired with a KUBTEC digital X-ray system in prone and lateral positions using consistent settings. Projected body area and lean area, defined as soft-tissue area excluding skeleton, were measured in ImageJ. The kyphosis index was measured in ImageJ on lateral radiographs as the perpendicular distance from the apex of the dorsal curvature to the line connecting the neck–back junction and tail base, divided by the length of that line. Three measurements were averaged. Excised femurs were scanned with a Quantum GX3 micro-CT system (Revvity). Cortical bone mineral density, thickness, area, and cortical-area fraction were quantified at a standardized mid-diaphyseal region spanning approximately 10% of femoral length. Ct.Ar/Tt.Ar was cortical area divided by total periosteal cross-sectional area. Analysis was performed blinded to treatment. Additional femoral measurements included bone volume and volume fraction, bone surface and surface-to-volume ratio, trabecular thickness, separation and number, structure model index, total and marrow cross-sectional areas, and growth-plate thickness.

### Single-cell RNA sequencing

Hearts and livers were collected at approximately 15 weeks of age from male and female CTL and mPSS mice. Each tissue comprised 12 libraries, with three biological replicates per sex-by-treatment group; each library was prepared from tissue pooled from three mice. Tissues were dissociated into single-cell suspensions according to the 10x Genomics tissue-dissociation protocol. Cell suspensions were filtered and assessed for concentration and viability before library preparation. Viable cells were processed with the 10x Genomics Single Cell 3′ v4 GEM-X workflow (PN-1000691) on Chromium X and sequenced with 150-bp paired-end reads.

### Reproductive and gonadal assessment

For reproductive assessment, mPSS-treated homozygous *Lmna^G^*^609^*^G/G^*^609^*^G^* males and females were pair-bred and monitored for mating, pregnancy, and delivery. Deliveries were recorded as vaginal or Caesarean following dystocia, and offspring were transferred to ICR foster dams. Offspring were genotyped by PCR, with a subset validated by Sanger sequencing across the *Lmna^G^*^609^*^G/G^*^609^*^G^* mPSS target region. At collection, gonads were excised, photographed, and processed for gross and histological analyses. Testes were weighed individually, and maximal ovary diameter was measured with a ruler in 0.5-mm increments. Bilateral testis weights and ovary diameters were averaged within each mouse before group comparison. For testis histomorphometry, three fields were averaged per mouse; ovarian endpoints were summarized per ovarian sample. Reserve follicles comprised primordial and primary follicles, identified by a single layer of flattened or cuboidal granulosa cells, respectively. Reserve follicle density was calculated as their combined count divided by the ovarian cross-sectional area (follicles/mm²). Gonadal H&E staining followed the protocol above.

### Analysis groups and nomenclature

For human fibroblast analyses, CTL denotes a non-targeting sgRNA. Bulk RNA-seq and ATAC-seq comprised unrelated WT, HG168, HG167 CTL-sgRNA, HG167 hPSS-sgRNA, WX CTL-sgRNA, and WX hPSS-sgRNA groups, with three biological replicates per group; CTL-sgRNA-treated WT and HG168 cells served as healthy references. For mouse single-cell analyses, CTL denotes PBS treatment. Human genomic analyses used hg38.

### Target-locus amplicon sequencing

Paired-end amplicon reads were merged with FLASH 1.2.11 using default overlap and mismatch settings. Reads containing an exact 157-nucleotide sequence spanning the wild-type *LMNA* c.1824C nucleotide and an unmodified hPSS target region were removed. The remaining reads were analyzed with CRISPResso2^5^ 2.1.3 against the HGPS-mutant amplicon using the 20-nucleotide hPSS guide sequence (5′-GGAGATGGGTCCACCCACCT-3′), a 10-nucleotide quantification window centred on the expected cleavage site, a 50-nucleotide display window, a minimum mean Phred quality score of 30, and a minimum single-base quality score of 10. Reads classified as unedited by CRISPResso2 were counted as unmodified, and all other classified reads were counted as modified. The mutant-specific modification fraction was calculated as modified reads divided by all reads remaining after removal of reads carrying the wild-type-specific sequence. Wild-type allele counts were obtained from the unmodified wild-type sequence rows in the allele tables before wild-type-sequence filtering and expressed as fractions of all aligned reads. The ten most abundant sequence classes were displayed individually.

### Off-target sequencing analysis

GUIDE-seq-2 libraries from HG167 and WX fibroblasts were analyzed using a workflow adapted from GUIDE-seq-2^2^ and TTISS^3^. The 8-nucleotide unique molecular identifier (UMI) at the 5′ end of read 1 was extracted with UMI-tools 1.1.0. Reads were trimmed with Cutadapt 4.7 using a minimum base quality of 20 and a minimum length of 25 nucleotides, aligned to hg38 with BWA-MEM2 2.2.1, and processed with SAMtools. Primary alignments with mapping quality ≥30 were retained. PCR duplicates were consolidated by paired-end directional UMI deduplication, and each deduplicated read pair was counted as one unique read.

DNA-integration endpoints were defined by the outer genomic coordinate of read 2. Endpoints separated by ≤10 bp were merged, and the coordinate supported by the most fragments represented the site. Candidate sites required at least two unique reads within a 20-nucleotide window. Reported site counts summed unique reads across each endpoint cluster; the maximum count in a 20-nucleotide window was reported separately. Sense and antisense profiles were calculated within a 60-nucleotide window centred on each site. Both genomic strands within 25 nucleotides of each site were searched for 20-nucleotide sequences adjacent to an `NGG` PAM and matching the hPSS guide sequence (5′-GGAGATGGGTCCACCCACCT-3′), allowing up to six substitutions. Sites were retained when the predicted Cas9 cleavage position was within 3 bp of the observed integration peak. Sites within 1 kb of hg38 `chr1:156138608` were assigned to the intended *LMNA* locus to account for the wild-type nucleotide in hg38.

The genomic context of these candidates was annotated against GENCODE v50 using ChIPseeker^6^ 1.38.0, and gene symbols were assigned with org.Hs.eg.db 3.18.0.

### WGS analysis of candidate sites

WGS reads were trimmed with Cutadapt^7^ 4.7 (minimum base quality, 20; minimum adapter overlap, 5 nucleotides; maximum error rate, 0.10; minimum length, 50 nucleotides), aligned to hg38 with BWA-MEM2 2.2.1, coordinate-sorted with SAMtools 1.9, and duplicate-marked with GATK^8^ 4.6.2.0/Picard 3.4.0. Alignment and mapping statistics were obtained with SAMtools `flagstat`, duplicate fractions with Picard `MarkDuplicates`, and mean coverage and coverage fractions with `CollectWgsMetrics`.

The *LMNA* target and the union of double-strand-break (DSB) profiling candidates from HG167 and WX with up to six guide-sequence mismatches were examined in both cell lines within ±200 bp of the expected cleavage site. Short variants were called with `Mutect2` by comparing each hPSS sample with its matched CTL sample, followed by orientation-bias modelling and `FilterMutectCalls`. `Mutect2` events were reported within ±200 bp, with cut-proximal filter status summarized within ±10 bp. Observed indel support was reported as x/N, where N comprised primary, nonduplicate fragments evaluable across the complete 21-bp cut-centred window and x comprised those carrying an indel within 10 bp of the cut.

Mapping and base quality thresholds were ≥20; overlapping mates were counted once. Indels required at least three high-quality aligned bases on each flank, and anchored deletions were allowed within the evaluable window. Incomplete or ambiguous indel evidence was excluded from x/N. The fragment-based evidence criterion was at least two indel-containing hPSS fragments and none in the matched CTL sample; observed counts were reported for all evaluable loci.

### Bulk RNA-seq processing

RNA-seq reads were aligned to hg38 with STAR^9^ 2.7.11b using an index generated from the GRCh38 primary assembly and GENCODE v49 annotation with `sjdbOverhang = 149`. Gene-level counts were generated from GENCODE v49 exon annotations with featureCounts^10^ 2.1.1 using paired-end, unstranded fragment counting. Lowly expressed genes were removed with `edgeR::filterByExpr`. Differential expression was analyzed with DESeq2^11^ Wald tests for HG167-hPSS versus HG167-CTL, WX-hPSS versus WX-CTL, and each CTL-sgRNA-treated HGPS line versus CTL-sgRNA-treated WT and HG168 cells. Differentially expressed genes were defined by a Benjamini–Hochberg-adjusted *P* value < 0.05 and an absolute log2 fold change ≥ log2(1.5).

For visualization, filtered counts were normalized by the trimmed mean of M values (TMM)^12^ method and transformed to log2 counts per million (logCPM). Sample correlations were calculated from the 5,000 most variable retained genes. Principal-component analysis used centred and scaled logCPM values for the union of genes differing between either HGPS line and both healthy references. The heatmap shows gene-wise Z scores of TMM-normalized logCPM values for genes classified as rescued-UP or rescued-DOWN in both HGPS lines.

### *LMNA* splice-junction analysis

Reads spanning the canonical *LMNA* exon 11–12 junction (`chr1:156138758–156139079`, plus strand) or the progerin-producing cryptic junction (`chr1:156138608–156139079`, plus strand) were obtained from STAR splice-junction output by summing uniquely and multiply mapping junction reads. Junction abundance was expressed as counts per million input reads. The progerin-junction fraction was calculated as cryptic-junction reads divided by the sum of cryptic- and canonical-junction reads. Replicate-level fractions were compared by Welch’s t-tests within two sets of three groups: HG168 CTL, HG167 CTL and HG167 hPSS; and WT CTL, WX CTL and WX hPSS. All three pairwise comparisons within each set were tested, with Benjamini–Hochberg correction across the six comparisons (*n* = 3 biological replicates per group).

### Expression rescue after hPSS

Disease-associated expression was defined separately in HG167 and WX. Genes significantly lower in CTL-treated HGPS cells than in both CTL-sgRNA-treated WT and HG168 cells were classified as disease-suppressed, whereas genes significantly higher than both healthy references were classified as disease-elevated. Each comparison was required to meet the differential-expression thresholds defined above.

Rescue sets were restricted to genes with annotated symbols. Disease-suppressed genes that increased significantly after hPSS treatment were classified as rescued-UP, and disease-elevated genes that decreased significantly were classified as rescued-DOWN. Shared rescued-UP and rescued-DOWN genes met the corresponding criteria in both HG167 and WX.

### Pathway enrichment analysis

Enrichment of biological-process and pathway annotations was tested separately for shared rescued-UP and rescued-DOWN genes and for genes significantly increased or decreased by hPSS in each fibroblast line. One-sided hypergeometric tests used all genes included in the corresponding DESeq2 analysis as background. Gene Ontology Biological Process and Kyoto Encyclopedia of Genes and Genomes (KEGG) enrichment were analyzed with clusterProfiler^14^ 4.10.1, Reactome enrichment with ReactomePA 1.46.0, and Molecular Signatures Database (MSigDB) Hallmark gene sets obtained with msigdbr 25.1.0 were analyzed with clusterProfiler.

For preranked gene-set enrichment analysis, all genes from each hPSS-versus-CTL comparison were ranked by the DESeq2 Wald statistic and analyzed with fgsea using 10,000 permutations. Benjamini–Hochberg correction was applied to over-representation and cell-line-wide GSEA analyses. Individual-pathway GSEA results in Supplementary Table 7 are reported as nominal P values.

### ATAC-seq processing

ATAC-seq adapters were trimmed with Cutadapt 4.6, retaining pairs in which both reads were at least 20 nt. Reads were aligned to hg38 with BWA-MEM2 2.2.1 and coordinate-sorted with SAMtools 1.6. PCR duplicates were removed with Picard `MarkDuplicates` (versions 3.1.1 and 3.4.0). Properly paired alignments with mapping quality ≥30 were retained; secondary, supplementary, and quality-control-failed alignments were excluded. Peaks were called independently in each library with MACS2^15^ 2.2.9.1 in paired-end mode at q < 0.01. Peak summits were expanded to 500-bp intervals, filtered against the hg38 blacklist, merged across libraries, recentred to 500 bp, and filtered against the blacklist again. A consensus peak was retained if it overlapped a peak called in at least two libraries within any of three categories: CTL-treated healthy-reference cells, CTL-treated HGPS cells, or hPSS-treated HGPS cells.

Paired-end fragments overlapping the consensus peaks were counted with featureCounts 2.0.6. Peaks with counts per million >1 in at least two libraries were analyzed with DESeq2 1.42.1 in R 4.3.3. Comparisons comprised hPSS-sgRNA versus CTL-sgRNA within HG167 and WX, and each CTL-treated HGPS line versus both CTL-treated healthy-reference lines, WT and HG168. Log2 fold changes were shrunken with ashr 2.2-63. Differentially accessible peaks were defined by a Benjamini–Hochberg-adjusted *P* value <0.05 and an absolute log2 fold change ≥log2(1.5). Quality-control analyses included mapping and chromosome-class read fractions, fragment-size distributions, fraction of reads in peaks, transcription-start-site enrichment, and Pearson correlations between samples. Correlations were calculated from TMM-normalized log2 counts per million for the 20,000 most variable consensus peaks, whereas principal-component analysis used TMM-normalized log2 counts per million for disease-associated peaks.

### Accessibility rescue after hPSS

Disease-associated accessibility was defined separately in HG167 and WX. Peaks significantly less accessible in CTL-treated HGPS cells than in both CTL-treated WT and HG168 cells were classified as disease-lost, whereas peaks significantly more accessible than both CTL-treated healthy references were classified as disease-gained. Both healthy-reference comparisons were required to meet the differential-accessibility thresholds. Disease-lost peaks that increased significantly after hPSS treatment were classified as rescued-open, and disease-gained peaks that decreased significantly were classified as rescued-closed. Shared rescued-open and rescued-closed peaks met the corresponding criteria in both HG167 and WX. Enrichment of hPSS-induced opening among disease-lost peaks and hPSS-induced closing among disease-gained peaks was tested using one-sided Fisher’s exact tests. For each cell line, the analysis included peaks tested in both comparisons with CTL-treated healthy references and the matched hPSS-versus-CTL comparison. *P* values were adjusted by the Benjamini–Hochberg method.

### ATAC-seq annotation and enrichment

ATAC-seq signal tracks normalized to reads per genomic content (RPGC) were generated with deepTools^16^ 3.5.4 from mapping-quality-filtered, properly paired, deduplicated alignments using 10-bp bins, the hg38 blacklist, and an effective genome size of 2,913,022,398; chrM and chrY were excluded from normalization. Signal heatmaps and aggregate profiles were calculated in 50-bp bins across 2-kb regions flanking peak centres, using the union of disease-lost or disease-gained peaks from HG167 and WX.

Peaks were annotated with ChIPseeker 1.38.0, `TxDb.Hsapiens.UCSC.hg38.knownGene` 3.18.0, and `org.Hs.eg.db` 3.18.0, with promoters defined as regions within 3 kb of a transcription start site. Gene Ontology Biological Process and Reactome enrichment were assessed for genes nearest to hPSS-increased and hPSS-decreased peaks in each cell line and to the HG167, WX, and shared rescued-open and rescued-closed peak sets. KEGG enrichment was assessed for genes nearest to the rescued peak sets. Known sequence motifs enriched among shared rescued-open and shared rescued-closed peaks were identified with HOMER^17^ 5.1 using the hg38 v7.0 annotation package and its default genomic background.

### RNA–ATAC integration

RNA-seq and ATAC-seq treatment effects were integrated separately for HG167 and WX. Peaks assigned to a gene within 3 kb of its transcription start site were considered promoter-linked. For genes linked to multiple promoter peaks, the median ATAC-seq log2 fold change for hPSS versus CTL was used. The analysis included disease-associated genes with finite RNA-seq and promoter-accessibility fold changes. A separate summary used all gene-linked peaks, with treatment effects again summarized by their median.

Rescued-UP genes were classified as concordant when both RNA expression and promoter accessibility increased after hPSS treatment; rescued-DOWN genes were concordant when both decreased. Promoter-linked peaks were not required to meet the differential-accessibility threshold. Representative loci were displayed using RPGC-normalized ATAC-seq tracks and RNA-seq counts-per-million summaries.

### Single-cell RNA-seq preprocessing

Heart and liver datasets were processed separately. Library was the unit used for quality control, count aggregation, and statistical comparison. Cell Ranger 9.0.0 generated feature–barcode matrices against GRCm39, including intronic reads.

Initial filtering was performed by library. Cells required ≥300 detected genes and ≥500 unique molecular identifiers (UMIs). Upper gene-count thresholds were set at the median plus three median absolute deviations (MADs), capped at 8,000 genes. Mitochondrial- and hemoglobin-transcript thresholds were set at the median plus three and five MADs, respectively, within ranges of 5–25% and 1–20%.

Doublets were identified with scDblFinder^18^ within each library and removed. Erythroid cells were identified from hemoglobin-transcript abundance and expression of *Hba-a1*, *Hba-a2*, *Hbb-bs*, *Hbb-bt*, *Alas2*, *Slc4a1*, *Klf1*, and *Gypa*. Erythroid-marker scores were calculated with Seurat `AddModuleScore` as mean log-normalized marker expression minus that of expression-matched control genes. Clusters containing ≥20 cells were removed when the median hemoglobin fraction was ≥2%, the median erythroid-marker score was ≥0.25, or ≥50% of cells contained ≥10% hemoglobin transcripts. Remaining cells with ≥10% hemoglobin transcripts were also removed.

Final heart cells required 500–6,000 detected genes, ≥1,000 UMIs, ≤10% mitochondrial transcripts, and ≤10% hemoglobin transcripts. Final liver cells required >500 and <5,000 detected genes, >1,000 and <40,000 UMIs, and <8% mitochondrial transcripts.

### Single-cell reduction and annotation

Counts were log-normalized with a scale factor of 10,000, and variable genes were selected using the variance-stabilizing transformation method. For heart, 3,000 variable genes were selected after excluding cell-cycle genes. Expression was scaled while regressing mitochondrial-transcript percentage, and principal-component analysis (PCA) was performed with 50 components. The first 20 components were used to construct the neighbor graph, generate the uniform manifold approximation and projection (UMAP), and cluster cells at resolution 0.5.

For liver, 3,000 variable genes were selected, and expression was scaled while regressing mitochondrial-transcript percentage. PCA was performed with 50 components, followed by Harmony correction for library-associated variation with `theta = 2`. The first 20 Harmony dimensions were used for the neighbor graph, UMAP, and clustering at resolution 0.4. Positive cluster markers were identified by Wilcoxon rank-sum tests, requiring expression in ≥20% of cells and log2 fold change ≥0.25, with up to 1,000 cells sampled per cluster. Cell types were assigned from canonical lineage markers. Liver annotation was additionally informed by an ambient-hepatocyte expression score calculated with `AddModuleScore` using *Alb*, *Ttr*, *Apoa1*, *Apoa2*, *Hnf4a*, *Otc*, *Bhmt*, and *Cps1*; an exocrine-contaminant cluster and a small cluster lacking a coherent lineage-marker profile were excluded.

### Cell composition and pseudobulk analysis

For each library, the abundance of each broad cell type was calculated as its fraction of all retained cells from the corresponding tissue. These values represent recovered-cell fractions. Cardiac fractions were analyzed with condition-only quasibinomial generalized linear models using the counts of the focal cell type and all remaining cells as the response; *P* values were adjusted across 13 cell types by the Benjamini–Hochberg method. Liver fractions were compared between CTL and mPSS libraries with two-sided Welch’s t tests followed by Benjamini–Hochberg correction across 14 cell types. Log2 ratios and absolute differences used pooled recovered-cell fractions within each treatment for heart and treatment-group mean library fractions for liver.

Pseudobulk counts were generated by summing raw counts within each cell type and library. For heart, cell type–library combinations containing <30 cells were excluded, and each analyzed cell type required at least two libraries per treatment. Genes required ≥10 total counts and ≥5 counts in at least two libraries. Differential expression was tested with DESeq2 using a treatment-only model with CTL as the reference. Differentially expressed genes had adjusted *P* < 0.05 and absolute log2 fold change ≥log2(1.5).

For liver, cell type–library combinations containing <20 cells were excluded; each cell type required ≥50 cells overall and at least two libraries per treatment. Genes required ≥10 total pseudobulk counts. DESeq2 used a sex-plus-treatment model, with a treatment-only model used when the sex-adjusted model could not be fitted. Differentially expressed genes had adjusted *P* < 0.05 and absolute log2 fold change ≥0.25.

For cardiac gene-set enrichment, genes were ranked by the DESeq2 Wald statistic. Liver genes were ranked by log2 fold change multiplied by −log10(adjusted *P*); log2 fold change alone was used when adjusted *P* values did not provide an informative ranking. Preranked enrichment analysis used mouse MSigDB Hallmark and Gene Ontology Biological Process gene sets containing 10–500 genes. Enrichment *P* values were adjusted by the Benjamini–Hochberg method. Normalized enrichment scores were oriented as mPSS relative to CTL. Pathways with false discovery rate (FDR) < 0.25 were eligible for display; positive hepatic normalized enrichment scores were displayed at zero.

### Cardiac fibroblast-state analysis

The 55,025 cardiac fibroblasts were log-normalized and reanalyzed using 3,000 variable genes. Mitochondrial- and hemoglobin-transcript percentages were regressed during scaling. PCA was performed with 50 components, and the first 15 components were used for the neighbor graph, UMAP, and clustering at resolution 0.15. Marker expression identified inflammatory/stress, extracellular matrix (ECM)-remodeling, and core structural fibroblast states.

Total fibroblasts and individual fibroblast states were quantified in each library as fractions of all retained heart cells. CTL and mPSS fractions were compared with two-sided Welch’s t tests followed by Benjamini–Hochberg correction across total fibroblasts and the three state measures. Sex-stratified values were displayed descriptively.

Fibroblast differential abundance was also analyzed with MiloR. A k-nearest-neighbor graph was constructed from PCs 1–15 with k = 20, and refined neighborhoods were sampled at a proportion of 0.10. Neighborhood cell counts were calculated by library and tested with a treatment-only model. Spatial FDR < 0.10 defined significant neighborhoods, which were assigned to the fibroblast state contributing the largest number of cells.

### Hepatic stromal-state analysis

Liver fibroblasts, hepatic stellate cells (HSCs), pericyte/vascular smooth-muscle cells (vSMCs), and mesothelial cells were extracted and reanalyzed using log-normalized expression, 2,500 variable genes, and PCA. The first 20 PCs were used for the neighbor graph and clustering at resolution 0.25. Clusters with non-stromal marker profiles were excluded. Marker expression identified ECM-remodeling fibroblast, HSC-like, fibroblast–HSC intermediate, mesothelial-like, pericyte/vSMC-contractile, and mural-myofibroblast-like states. The retained 3,333 cells were embedded for display using the first 15 PCs.

Each stromal state was quantified per library as a fraction of all retained liver cells. CTL and mPSS fractions were compared with two-sided Welch’s t tests followed by Benjamini–Hochberg correction across the six states. The remodeling-state composite was defined as the sum of the fibroblast–HSC intermediate, pericyte/vSMC-contractile, and mural-myofibroblast-like fractions and was tested separately with a two-sided Welch’s t test.

Expression analysis was restricted to the three states with sufficient cells in both treatments: ECM-remodeling fibroblasts, HSC-like cells, and mesothelial-like cells. Library-level mean expression of *Lrat*, *Rbp1*, *Dcn*, *Tgfbi*, *Mif*, *Cxcl12*, *Icam1*, and *Col1a1* was compared by two-sided Welch’s t tests, followed by Benjamini–Hochberg correction across the 24 gene–state combinations. Structural-ECM heatmaps used broad-cell-type pseudobulk DESeq2 fold changes and adjusted *P* values.

### Cell–cell communication

Cell–cell communication was inferred separately for CTL and mPSS cells using CellChat 2.2.0.9001 and CellChatDB.mouse. Communication probabilities were estimated using a 10% trimmed mean without weighting by cell-group abundance, followed by pathway-level network and centrality analysis. Cells were pooled within each treatment; communication changes were therefore descriptive and were not subjected to library-level hypothesis testing. For heart, cell groups and interactions required at least 20 cells. The pathway summary included MIF, CCL, CXCL, TNF, ICAM, ANNEXIN, CypA, SIRP, IL1, and SPP1 signaling. The displayed immune- and vascular-to-fibroblast interactions had a maximum communication probability >0.005 and lower probabilities after mPSS treatment. Interactions involving proliferating-cell groups or the broadly distributed *Ppia*–*Bsg* axis were excluded.

For liver, cell groups required ≥20 cells in each treatment, and interactions required a minimum group size of 10 cells. T and natural killer (NK) cells were combined, and up to 800 cells were sampled per cell group and treatment. The pathway summary included MIF, CCL, CXCL, TNF, IL1, SPP1, THBS, SIRP, ICAM, VCAM, and SEMA4 signaling.

Liver communication analysis used monocyte-derived macrophages, Kupffer macrophages, neutrophils, and T/NK cells as senders and ECM-remodeling fibroblast, HSC-like, and mesothelial-like states as receivers. The evaluated ligand–receptor pairs comprised *Mif*–*Ackr3*, *Tnf*–*Tnfrsf1a/b*, *Ccl5*–*Ackr1/2*, *Sema4a*–(*Nrp1*+*Plxna1*) or *Plxnb2*, *Sirpb1b*–*Cd47*, *Thbs1*–*Cd47* or *Sdc4*, *Vcam1*–(*Itga4*+*Itgb1*) or (*Itga4*+*Itgb7*), and *Spp1*–(*Itga5*+*Itgb1*) or (*Itgav*+*Itgb1*). The 91 displayed interactions comprised all sender–receiver–ligand–receptor combinations among these groups and pairs with a CellChat-reported P value < 0.05 in either treatment. Expression of *Cxcl2*, *Mif*, and *Tnf* was summarized as the mean log-normalized expression within each library for monocyte-derived macrophages, Kupffer macrophages, and neutrophils.

### Single-cell software

Single-cell analyses used R 4.3.3, Seurat^19^ 5.2.1, and SeuratObject 5.0.2. Doublet detection used scDblFinder^18^, liver integration used Harmony^20^, and pseudobulk differential expression used DESeq2 1.42.1. Cardiac enrichment used clusterProfiler; liver enrichment used fgsea 1.28.0 with gene sets obtained through msigdbr 25.1.0. Neighborhood and communication analyses used MiloR 2.9.1 and CellChat 2.2.0.9001, respectively. Random seeds were 1234 for preprocessing and pseudobulk workflows and 20260702 for liver CellChat analysis.

### Statistical analysis

Statistical analyses were performed in GraphPad Prism 10.2.3.347 and R 4.3.3. Unless stated otherwise, tests were two-sided. Survival was estimated by the Kaplan–Meier method and compared with the log-rank (Mantel–Cox) test. Two-group endpoints were compared with unpaired Student’s *t* tests or Welch’s *t* tests as specified in the figure legends. One- or two-way ANOVA with Tukey correction was used for the multiple-group comparisons specified in the figure legends. Benjamini–Hochberg correction was applied to the comparison sets specified above. For the three-protein comparisons specified in the figure legends, unpaired Student’s t-tests used two-stage Benjamini–Krieger–Yekutieli correction across the three proteins within each cell model at a target FDR of 1%. Corrected results are reported as adjusted *P* values or *q* values, as indicated in the figure legends. Data are presented as mean ± SEM unless stated otherwise. *P* values, sample sizes, and analysis units are reported in the Results, figure legends, and source data. *P* < 0.05 was considered significant; adjusted *P* or *q* values were used where multiple-testing correction was applied, unless an analysis-specific FDR threshold is stated.

