## Extended Data Figures 1-9 for "Genomic splice-donor disruption rescues systemic disease in progeria"

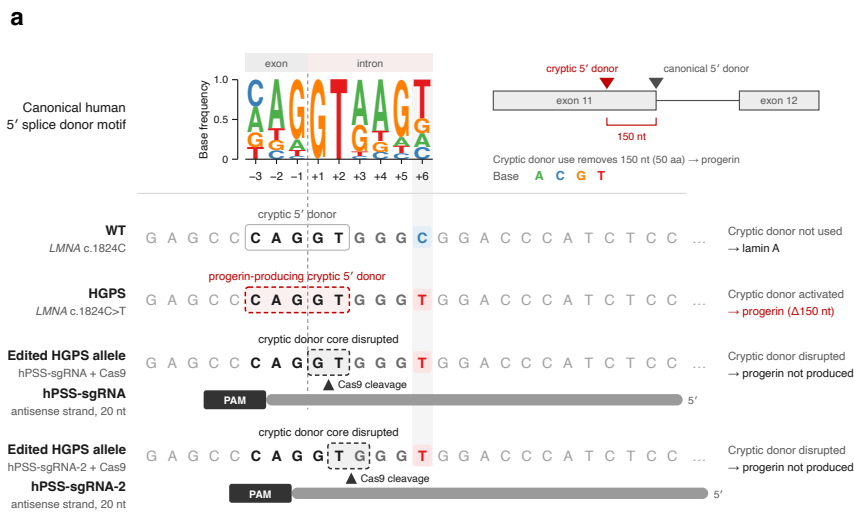

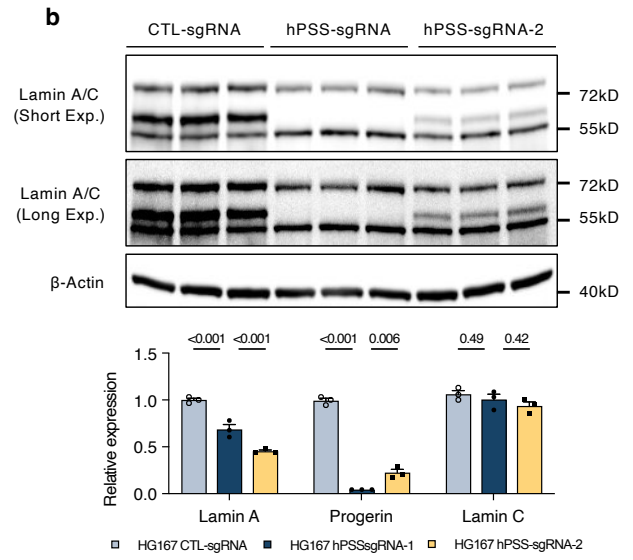

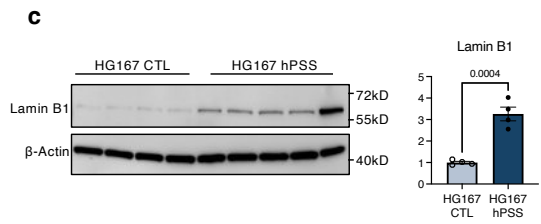

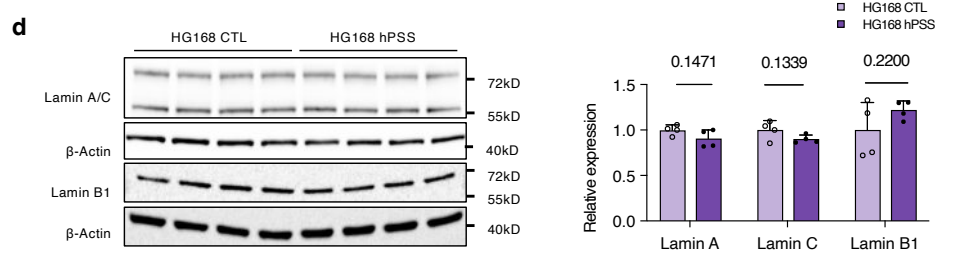

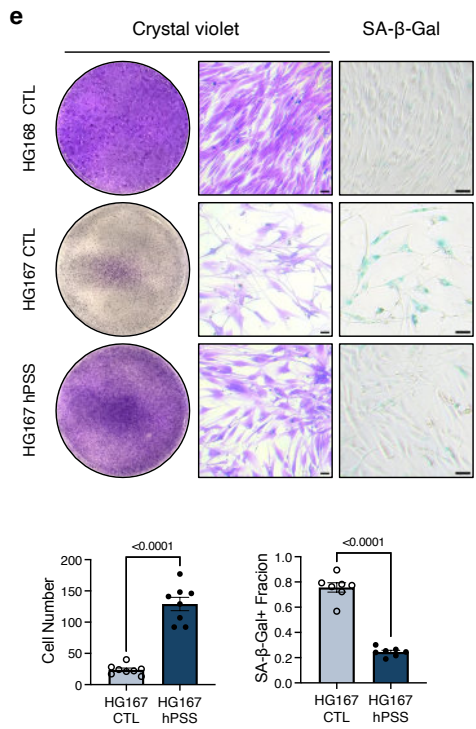

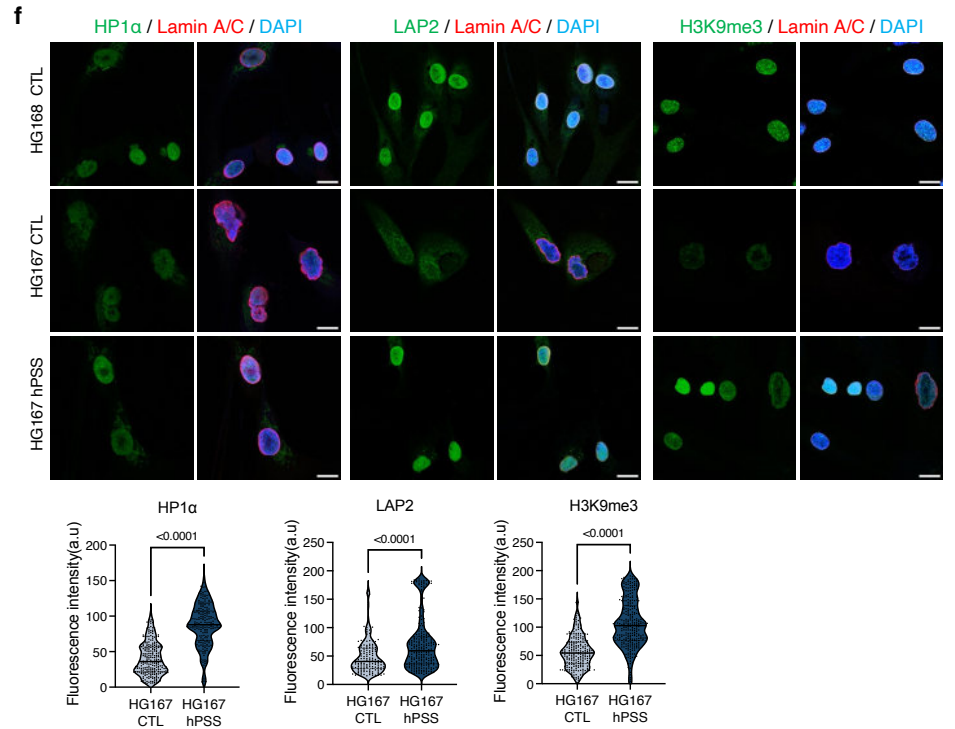

**Extended Data Fig. 1 | Guide-offset comparison and cellular responses to hPSS in human fibroblasts.**

**a**, Cryptic-donor targeting by human progerin splice-site silencing sgRNA (hPSS) and its one-nucleotide-offset guide, hPSS-sgRNA-2. **b**, Lamin A/C immunoblots at short and long exposures in HG167 HGPS fibroblasts treated with non-targeting sgRNA (CTL), hPSS or hPSS-sgRNA-2, with protein abundance normalized to β-Actin and CTL ($n = 3$ biological replicates per group). **c,d**, Lamin B1 in HG167 (c) and lamin A, lamin C and lamin B1 in healthy-reference HG168 fibroblasts (d), normalized to β-Actin and matched CTL ($n = 4$ independent cultures per group). **e**, Crystal-violet and senescence-associated β-galactosidase (SA-β-gal) staining; cell counts and SA-β-gal-positive fractions ($n = 8$ and $7$ fields per group, respectively, from three independent cultures). Scale bars, $100\ \mu\text{m}$. **f**, Heterochromatin protein 1α (HP1α), lamina-associated polypeptide 2 (LAP2) and histone H3 lysine 9 trimethylation (H3K9me3) staining. Violin plots show single-cell intensities from three independent cultures (CTL/hPSS: HP1α, 253/204; LAP2, 150/244; H3K9me3, 237/283 cells). Scale bars, $10\ \mu\text{m}$. CTL-treated HG168 fibroblasts provide the healthy reference. Bars show mean ± s.e.m. Comparisons used Student's $t$-tests (c,d), Welch's $t$-tests (e,f), or two-way ANOVA with Tukey correction (b).

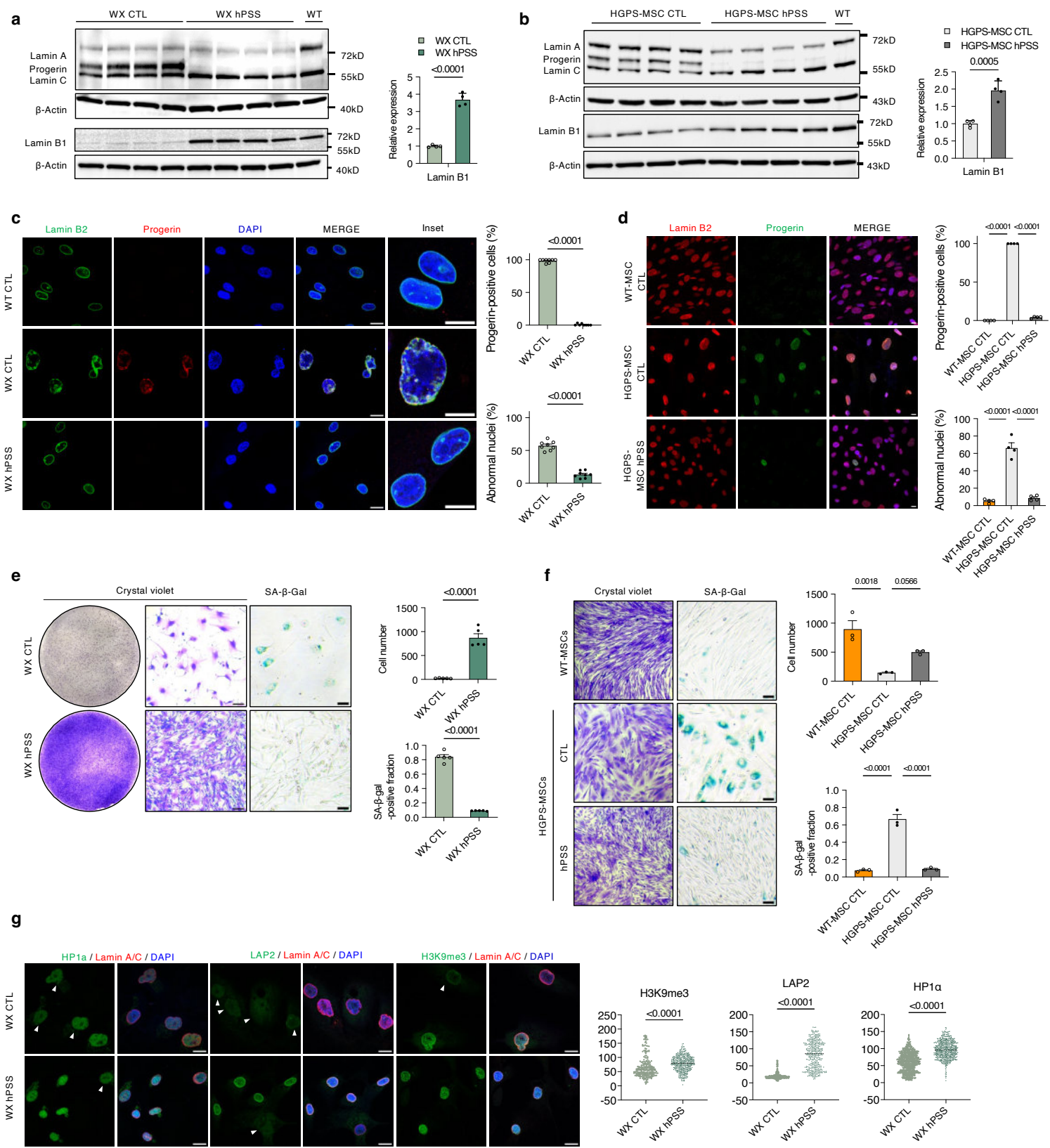

**Extended Data Fig. 2 | Cellular responses to hPSS in WX HGPS fibroblasts and HGPS-derived mesenchymal stem cells.**

**a,b**, Lamin A/C, progerin and lamin B1 immunoblots in WX HGPS fibroblasts (a) and HGPS induced pluripotent stem cell-derived mesenchymal stem cells (MSCs; b) treated with non-targeting sgRNA (CTL) or human progerin splice-site silencing sgRNA (hPSS), with lamin B1 normalized to β-actin and matched CTL ($n = 4$ biological replicates per group). **c,d**, Lamin B2, progerin and DAPI staining, with progerin-positive cells and abnormal nuclei in WX (c; $n = 8$ fields per group) and MSCs (d; $n = 4$ measurements per group). Scale bars, 10 μm. **e,f**, Crystal-violet and senescence-associated β-galactosidase (SA-β-gal) staining, cell counts and SA-β-gal-positive fractions in WX (e; $n = 5$ measurements per group) and MSCs (f; $n = 3$ per group). Scale bars, 100 μm. **g**, Heterochromatin protein 1α (HP1α), lamina-associated polypeptide 2 (LAP2) and histone H3 lysine 9 trimethylation (H3K9me3) staining and fluorescence intensities (CTL/hPSS: 626/592, 402/375 and 240/439 cells, respectively) in CTL- and hPSS-treated WX HGPS fibroblasts. Scale bars, 10 μm. Wild-type (WT) fibroblasts and MSCs received CTL and served as healthy references. Bars show mean ± s.e.m. Comparisons used unpaired Student's *t*-tests (a–c,e,g) or one-way ANOVA with Tukey correction (d,f).

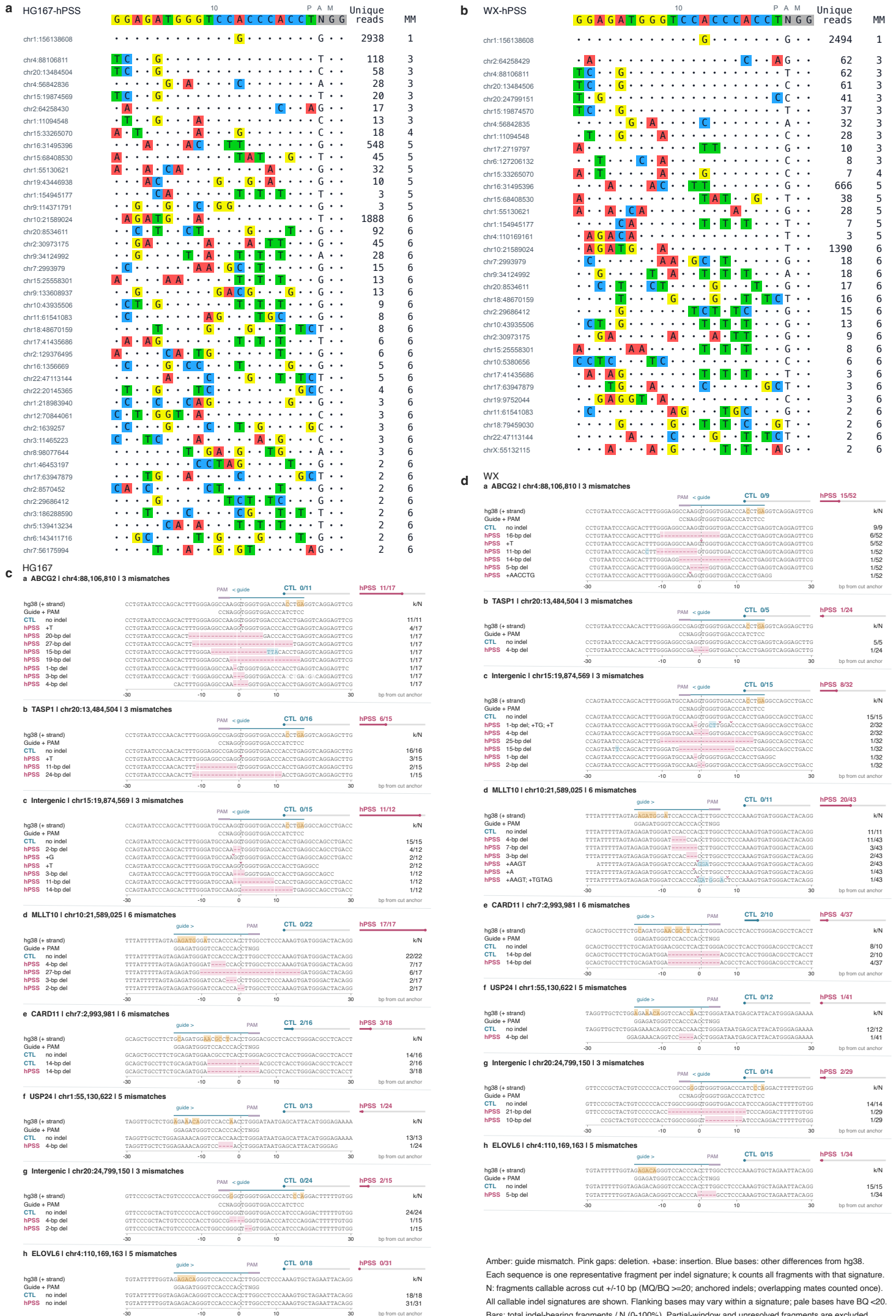

Amber: guide mismatch. Pink gaps: deletion. +base: insertion. Blue bases: other differences from hg38.

Each sequence is one representative fragment per indel signature; k counts all fragments with that signature.

N: fragments callable across cut +/-10 bp (MQ/BQ >=20); anchored indels; overlapping mates counted once.

All callable indel signatures are shown. Flanking bases may vary within a signature; pale bases have BQ <20.

Bars: total indel-bearing fragments / N (0-100%). Partial-window and unresolved fragments are excluded.

**Extended Data Fig. 3 | GUIDE-seq2 and whole-genome sequencing of hPSS candidate off-target sites.**

**a,b**, GUIDE-seq2 sites in HG167 (a; 42 sites) and WX (b; 33 sites) HGPS fibroblasts treated with human progerin splice-site silencing sgRNA (hPSS), including the on-target site and all candidates with up to six guide mismatches. Coordinates indicate integration peaks in hg38. Dots denote matches to the guide; coloured bases denote mismatches. PAM, protospacer-adjacent motif; MM, protospacer substitutions excluding the PAM; Unique reads, unique molecular identifier (UMI)-deduplicated read pairs supporting each site. The on-target mismatch reflects the hg38 wild-type *LMNA* sequence. **c,d**, Representative whole-genome sequencing (WGS) fragment alignments at eight candidate loci in HG167 (c) and WX (d), comparing hPSS with matched non-targeting sgRNA controls (CTL).

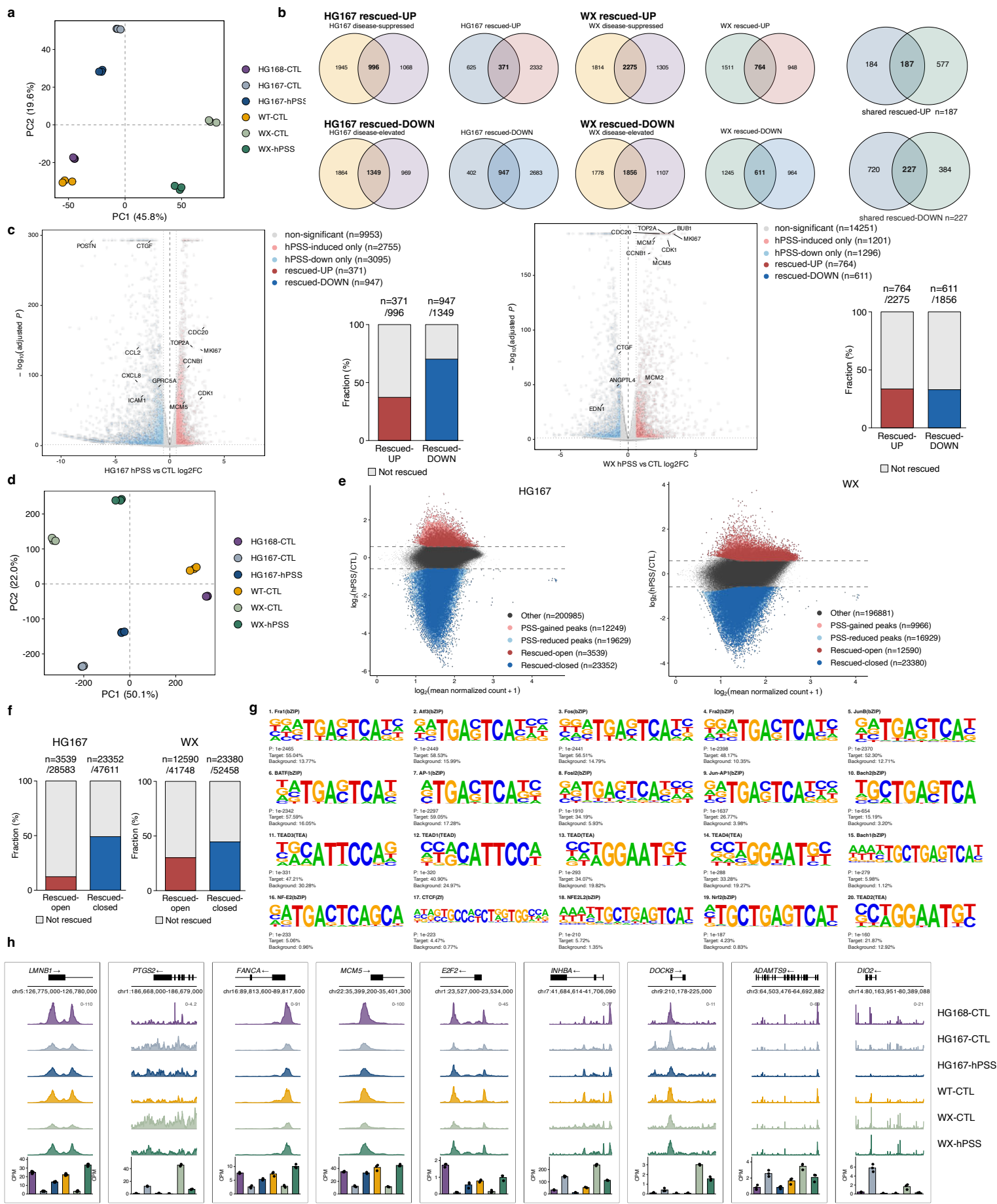

##### **Extended Data Fig. 4 | Transcriptional and chromatin-accessibility responses to hPSS.**

**a**, Principal-component analysis of disease-associated RNA expression in HG167 and WX HGPS fibroblasts treated with human progerin splice-site silencing sgRNA (hPSS) or non-targeting sgRNA (CTL), with CTL-treated wild-type (WT) and HG168 fibroblasts as healthy references. **b**, Gene-set intersections. Disease-suppressed/elevated genes differed in the same direction from both healthy references; rescued-UP/DOWN genes additionally changed in the opposite direction after hPSS (see Method). **c**, Treatment volcano plots and fractions of disease-associated genes rescued by hPSS. **d**, Principal-component analysis of disease-associated ATAC-seq peaks. **e**, Treatment MA plots. **f**, Rescued-peak fractions. Disease-lost/gained peaks had lower/higher accessibility than both healthy references; rescued-open/closed peaks changed oppositely after hPSS. All required contrasts met Benjamini–Hochberg-adjusted $P < 0.05$ and $\geq 1.5$-fold change (DESeq2 Wald tests). **g**, Twenty known sequence motifs enriched among shared rescued-closed peaks (HOMER). **h**, ATAC-seq tracks and RNA abundance at the indicated loci. Tracks show reads per genomic content; bars show trimmed mean of M values (TMM)-normalized counts per million (mean ± s.e.m.; points, biological replicates). $n = 3$ biological replicates per group.

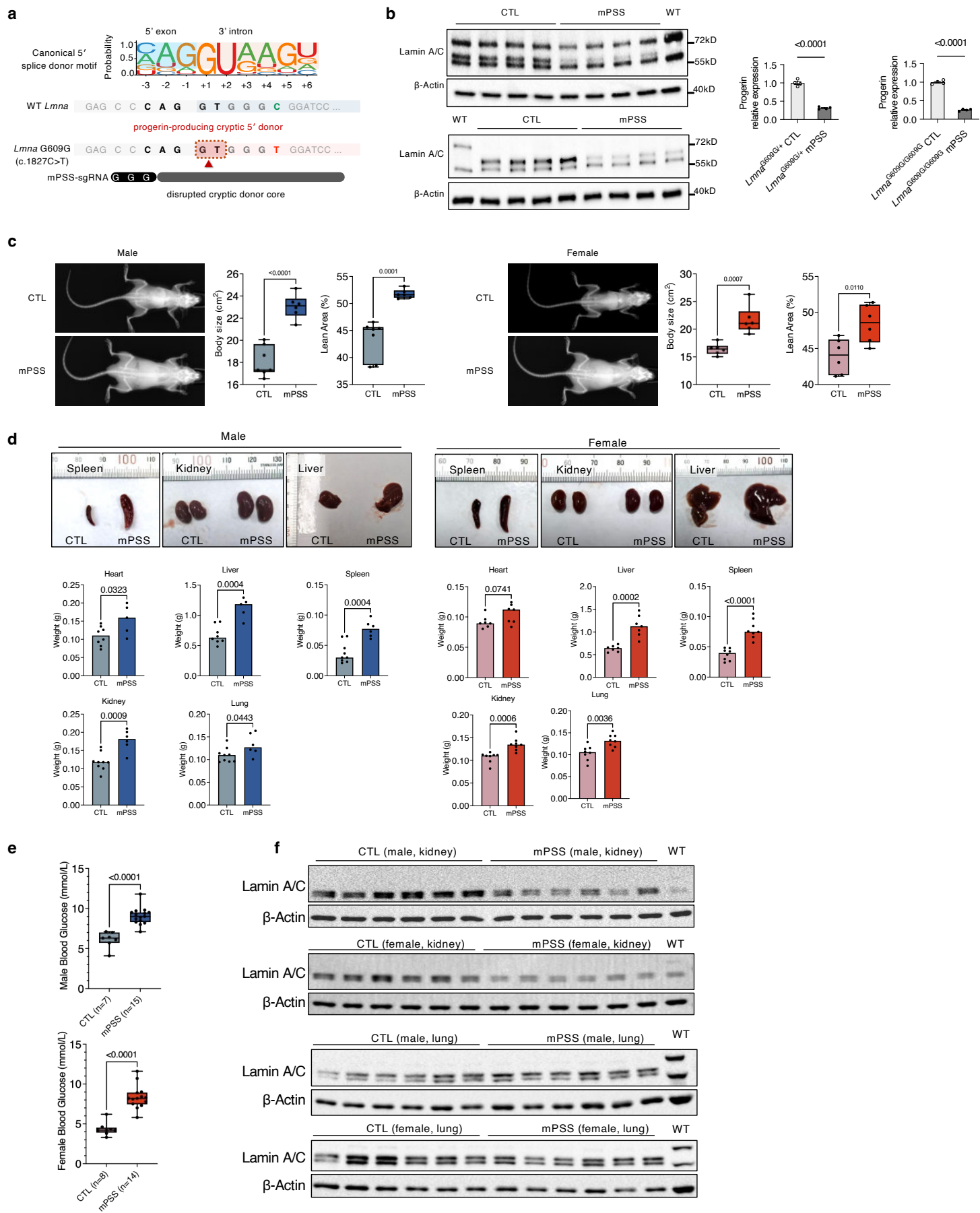

**Extended Data Fig. 5 | Mouse-target validation and systemic phenotypes after mPSS treatment.**

**a**, Design of *Lmna* cryptic-donor targeting by mouse progerin splice-site silencing sgRNA (mPSS). **b**, Lamin A/C immunoblots and progerin abundance in *Lmna*<sup>G609G/+</sup> and *Lmna*<sup>G609G/G609G</sup> mouse embryonic fibroblasts (MEFs) receiving non-targeting sgRNA (CTL) or mPSS-sgRNA, normalized to β-Actin and CTL ($n = 4$ biological replicates per group). Wild-type (WT) MEFs serve as a reference. **c**, Radiographs, body area and lean-area percentage in Cas9-expressing *Lmna*<sup>G609G/G609G</sup> mice receiving PBS (CTL) or AAV9 carrying mPSS-sgRNA (CTL/mPSS: male, $n = 7/6$; female, $n = 6/6$ mice). **d**, Organ specimens and weights. Male CTL/mPSS: heart/liver, $n = 8/5$; spleen/lung/kidney, $n = 9/6$ mice. Female: heart/liver, $n = 7/7$; spleen/lung/kidney, $n = 8/8$ mice. Kidney values are bilateral means. **e**, Random-fed blood glucose (male, $n = 7/15$; female, $n = 8/14$ mice). **f**, Kidney and lung lamin A/C and progerin immunoblots, with β-actin loading controls and WT references ($n = 6$ tissue samples per sex, tissue and treatment). Bars show mean ± s.e.m.; boxes show medians, interquartile ranges and minimum-to-maximum whiskers. Points denote replicates (b) or mice (c–e). Comparisons used unpaired Student's *t*-tests (b–d) or Welch's *t*-tests (e).

**a**

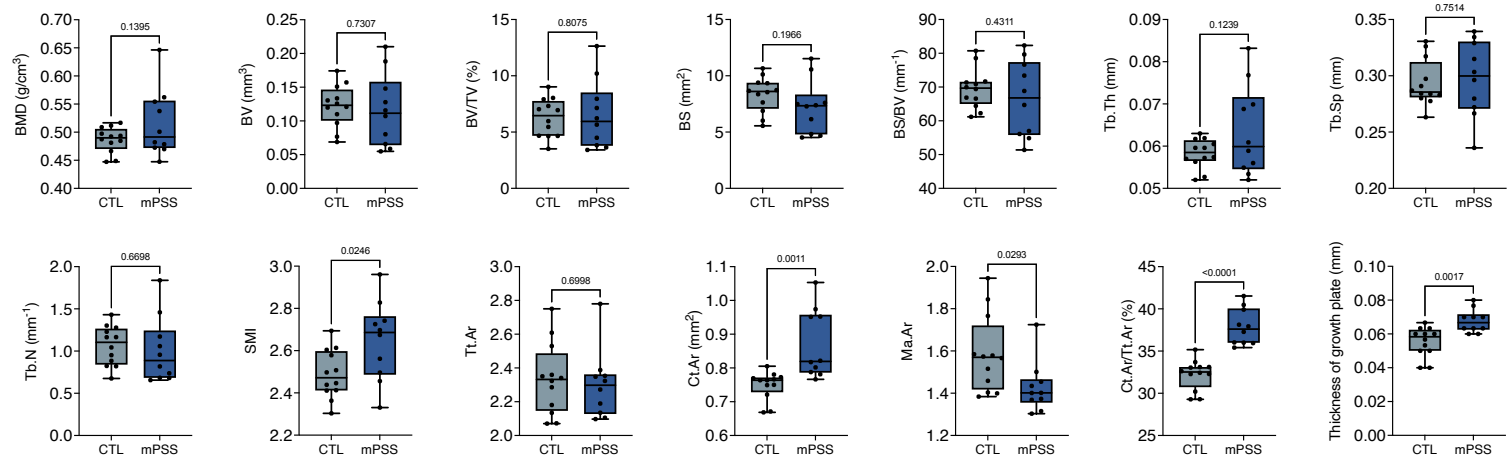

**b**

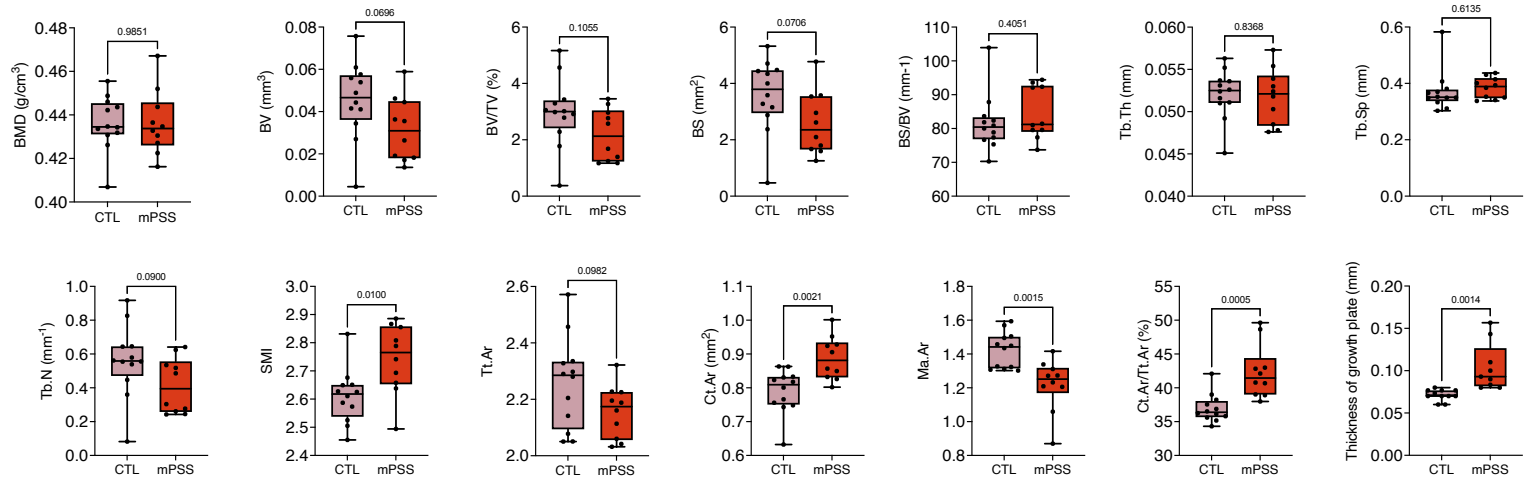

**c**

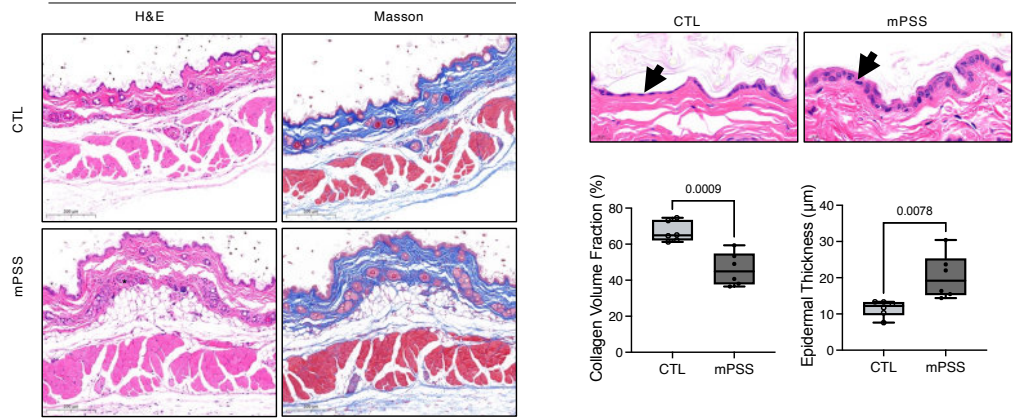

**Extended Data Fig. 6 | Additional skeletal and skin histopathological analyses after mPSS treatment.**

**a,b,** Femoral microcomputed tomography measurements in male (a) and female (b) *Lmna*<sup>G609G/G609G</sup> mice receiving PBS (CTL) or AAV9 carrying mouse progerin splice-site silencing sgRNA (mPSS): bone mineral density (BMD), bone volume (BV), bone volume fraction (BV/TV), bone surface (BS), surface-to-volume ratio (BS/BV), trabecular thickness (Tb.Th), separation (Tb.Sp) and number (Tb.N), structure model index (SMI), total (Tt.Ar), cortical (Ct.Ar) and marrow (Ma.Ar) cross-sectional areas, cortical area fraction (Ct.Ar/Tt.Ar), and growth-plate thickness. CTL, $n = 12$; mPSS, $n = 10$ femora per sex; female growth-plate measurements, $n = 11$ and 9, respectively. **c,** Skin haematoxylin and eosin (H&E) and Masson's trichrome staining, collagen volume fraction and epidermal thickness ($n = 6$ sex-pooled samples per group). Arrows indicate the epidermis. Scale bars, 200 $\mu\text{m}$. Boxes show medians and interquartile ranges, with minimum-to-maximum whiskers. Comparisons used unpaired Student's $t$-tests.

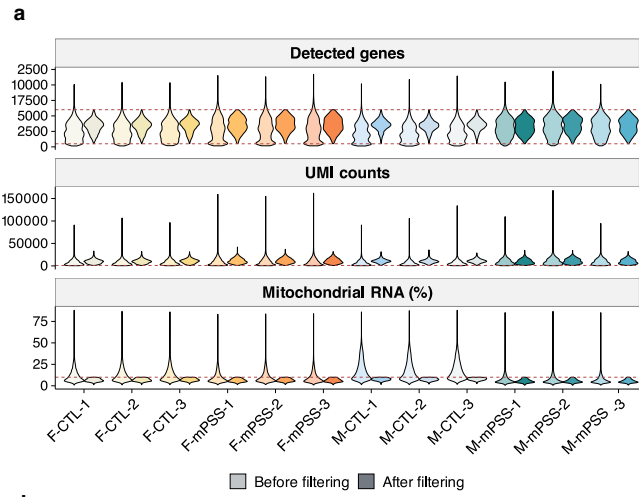

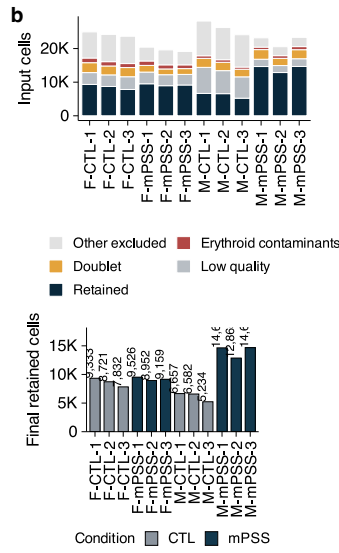

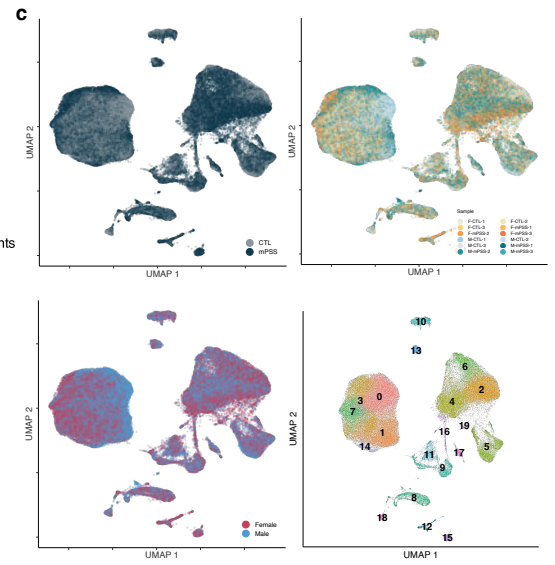

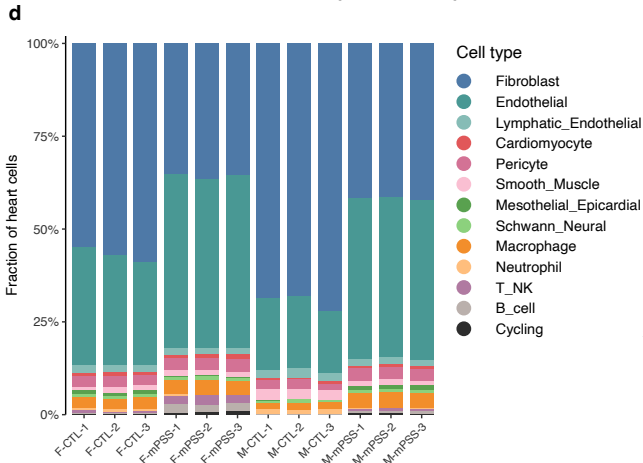

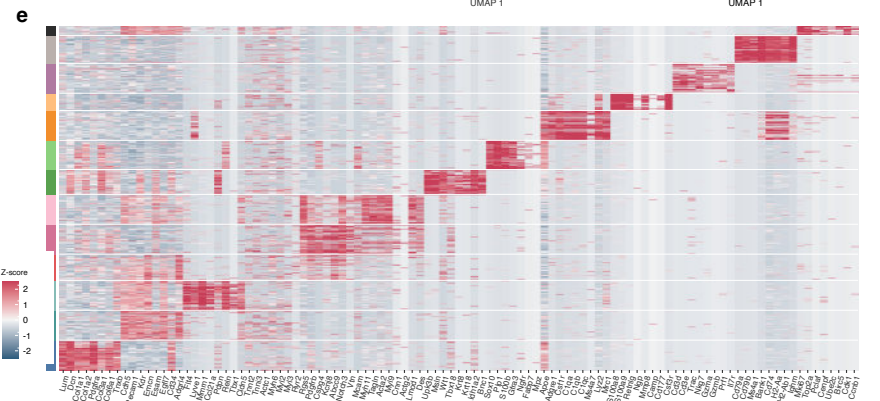

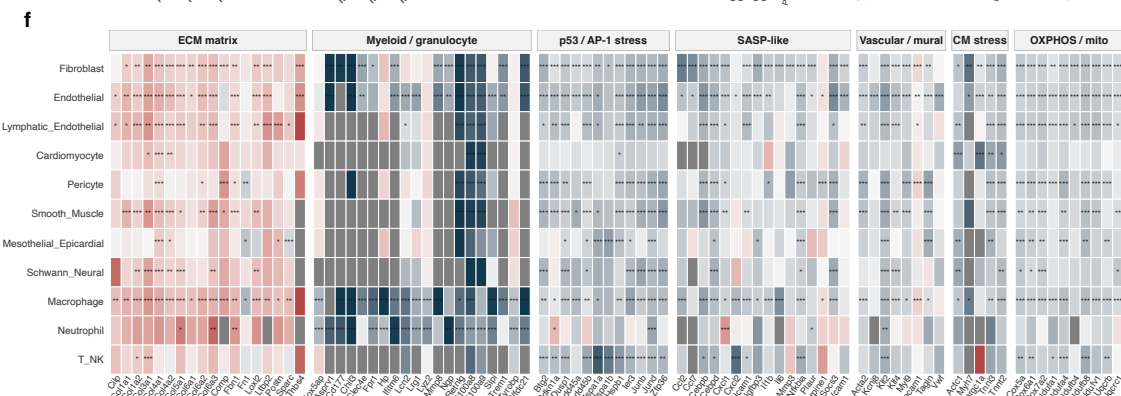

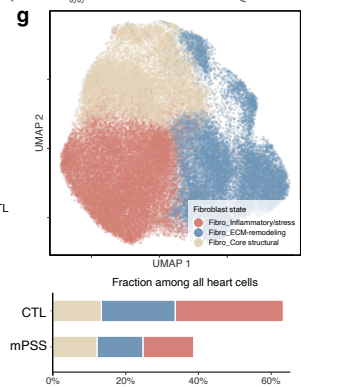

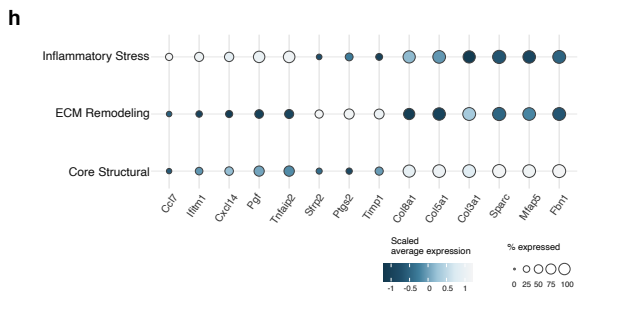

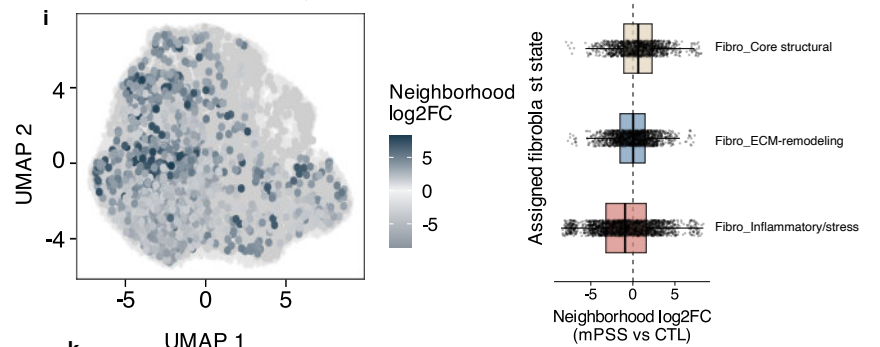

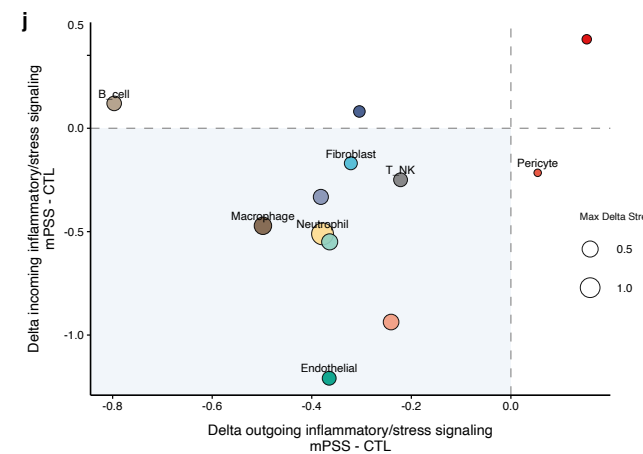

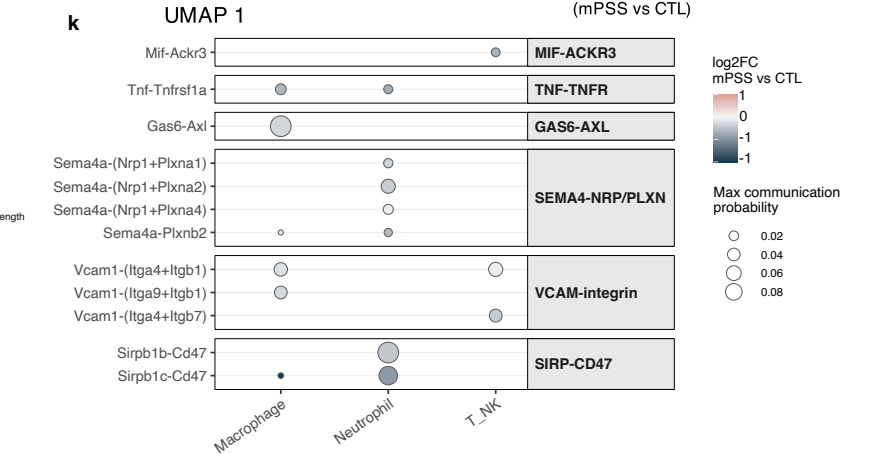

**Extended Data Fig. 7 | Heart single-cell RNA-seq quality control and fibroblast-state analyses.**

**a**, Detected genes, unique molecular identifier (UMI) counts and mitochondrial RNA before/after filtering of heart cells from *Lmna*<sup>G609G/G609G</sup> mice treated with PBS (CTL) or AAV9 carrying mouse progerin splice-site silencing sgRNA (mPSS); dashed thresholds indicate 500–6,000 genes, $\geq 1,000$ UMIs and $\leq 10\%$ mitochondrial RNA. **b**, Filtering disposition and retained-cell counts. **c**, UMAPs coloured by treatment, library, sex and cluster. **d**, Cell-type fractions per library. **e**, Scaled cell-type marker expression. **f**, Cardiomyocyte (CM), oxidative phosphorylation (OXPHOS)/mitochondrial (mito), senescence-associated secretory phenotype (SASP) and extracellular matrix (ECM) programmes: pseudobulk gene-expression $\log_2$ fold changes (mPSS/CTL; DESeq2 Wald tests, Benjamini–Hochberg correction; \*, \*\*, \*\*\* denote adjusted $P < 0.05, 0.01, 0.001$). **g**, Fibroblast-state UMAP and fractions among heart cells. **h**, Fibroblast-state marker expression. **i**, MiloR neighbourhood abundance changes; colour identifies spatial FDR $< 0.10$ and indicates $\log_2$ fold change. Boxes show medians and interquartile ranges (IQRs), with whiskers within 1.5 IQRs; points denote neighbourhoods. **j**, CellChat-inferred incoming and outgoing signalling changes. **k**, Inferred immune-to-fibroblast ligand–receptor communication. T\_NK, T/natural killer cells. The heart dataset comprised 12 libraries, with three biological replicates per sex-by-treatment group.

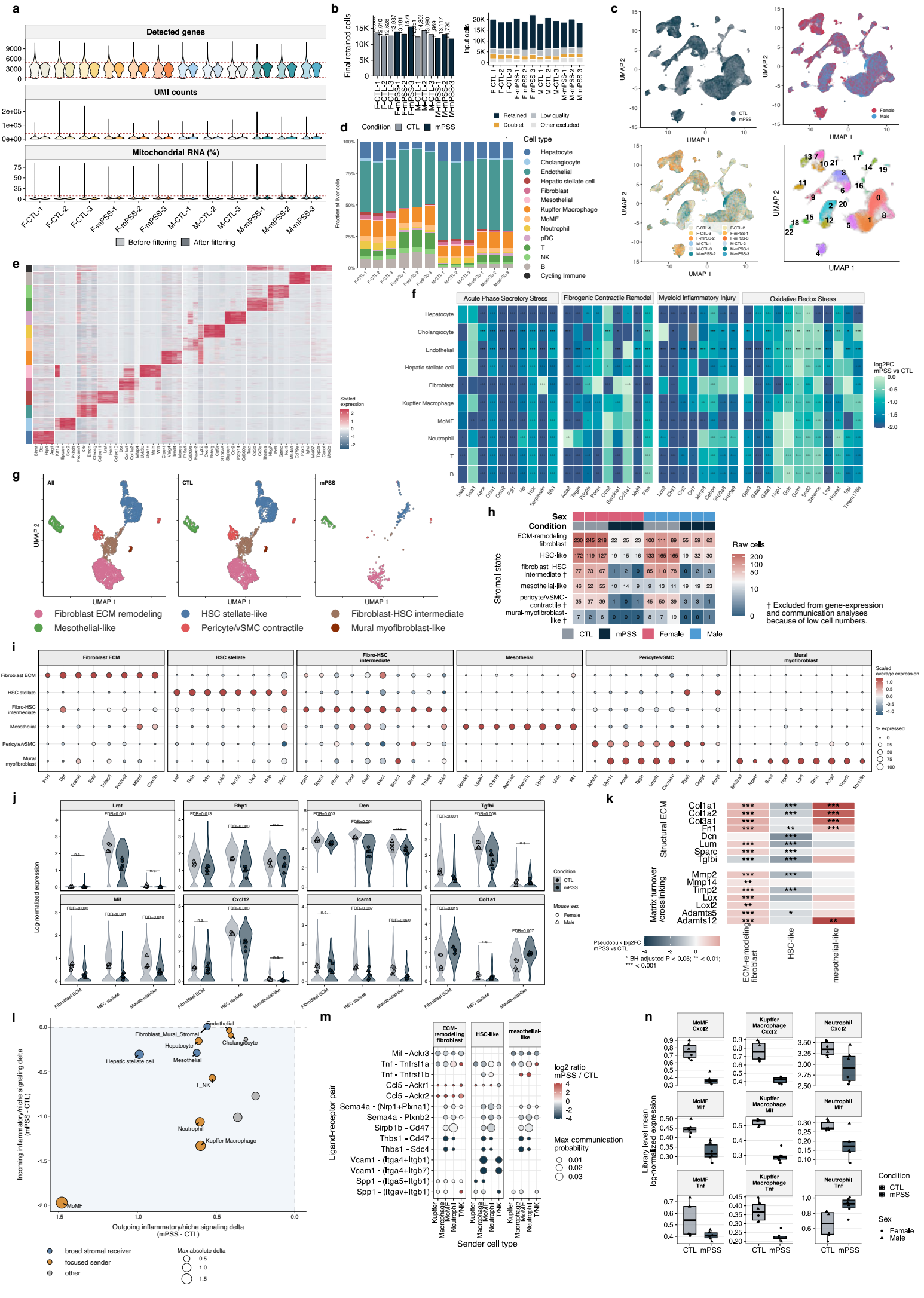

### Extended Data Fig. 8 | Liver single-cell RNA-seq quality control and stromal-immune analyses.

**a,b**, Liver single-cell RNA-seq quality-control distributions from *Lmna*<sup>G609G/G609G</sup> mice treated with PBS (CTL) or AAV9 carrying mouse progerin splice-site silencing sgRNA (mPSS) (a; thresholds: >500 and <5,000 genes, >1,000 and <40,000 unique molecular identifiers, <8% mitochondrial RNA), filtering disposition and cell yields (b). **c**, UMAPs by treatment, sex, library and cluster. **d,e**, Cell-type fractions (d) and scaled marker expression (e). **f**, Pseudobulk gene-expression log<sub>2</sub> fold changes (mPSS/CTL). **g–i**, Stromal-state UMAPs (g), cell counts per library (h) and marker expression (i). HSC, hepatic stellate cell; vSMC, vascular smooth muscle cell. **j**, Single-cell expression distributions with library-level means; Welch's *t*-tests with Benjamini–Hochberg correction across 24 gene–state comparisons (*n* = 6 libraries per treatment); n.s., adjusted *P* ≥ 0.05. **k**, Extracellular matrix (ECM) gene-expression changes in fibroblasts, hepatic stellate cells and mesothelial cells. In **f,k**, DESeq2 Wald tests used Benjamini–Hochberg correction; \*, \*\*, \*\*\* denote adjusted *P* < 0.05, 0.01, 0.001. **l**, CellChat-inferred signalling changes. **m**, Ninety-one inferred immune-to-stromal ligand–receptor interactions. MoMF, monocyte-derived macrophages; TNK, T/natural killer cells. **n**, Library-mean *Cxcl2*, *Mif* and *Tnf* expression. Points in n denote libraries (*n* = 6 per treatment). Boxes show medians and interquartile ranges (IQRs), whiskers within 1.5 IQRs. The liver dataset comprised 12 libraries, with three biological replicates per sex-by-treatment group.

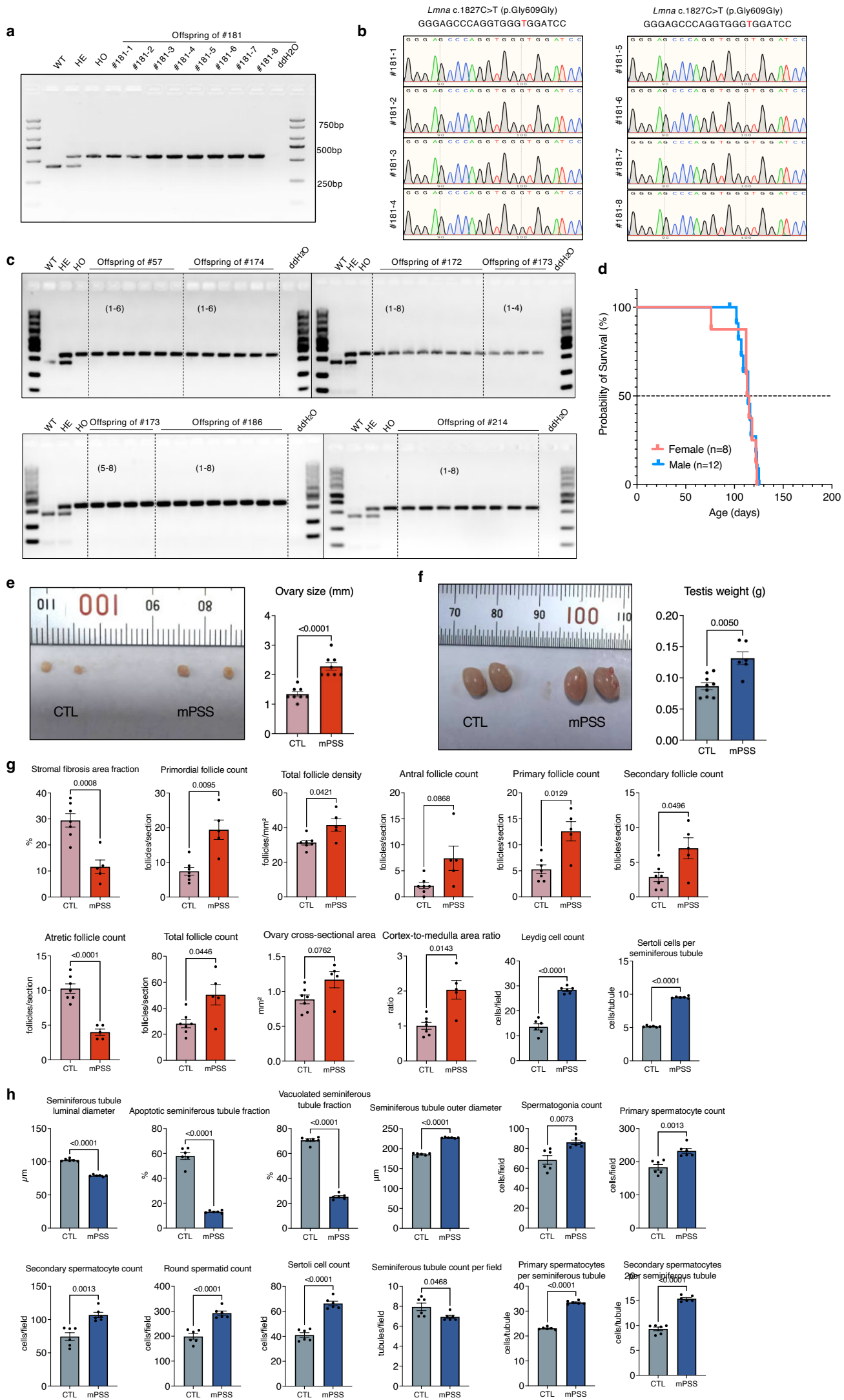

#### Extended Data Fig. 9 | Offspring genotyping and additional gonadal histomorphometry.

**a**, Genotyping PCR for offspring #181-1–181-8, with wild-type (WT), heterozygous (HE), homozygous (HO) and water controls. **b**, Sanger traces across the *Lmna* G609G locus in these eight offspring. **c**, PCR genotyping of the remaining 44 offspring. **d**, Kaplan–Meier survival of *Lmna*<sup>G609G/G609G</sup> offspring (female, $n = 8$; male, $n = 12$; pooled median, 114 days); ticks indicate censoring. **e,f**, Gross ovaries and testes from *Lmna*<sup>G609G/G609G</sup> mice treated with PBS (CTL) or mouse progerin splice-site silencing sgRNA (mPSS), with bilateral mean ovary size (**e**; $n = 8$ biological replicates per group) and testis weight (**f**; CTL/mPSS, $n = 9/6$ biological replicates). **g**, Ovarian stromal fibrosis, follicle counts/density, cross-sectional area and cortex-to-medulla ratio (CTL/mPSS, $n = 7/5$ ovarian samples), with Leydig cells per field and Sertoli cells per seminiferous tubule ($n = 6$ mice per group). **h**, Seminiferous-tubule diameters, apoptotic/vacuolated fractions, germ-cell and Sertoli-cell counts, tubules per field and spermatocytes per tubule. Testicular measurements average three fields per mouse ($n = 6$ per group). Data are mean ± s.e.m. Comparisons in **e–h** used Welch's $t$-tests.
